# Global gene-expression analysis reveals the molecular processes underlying ClC-5 loss-of-function in novel Dent Disease 1 cellular models

**DOI:** 10.1101/2020.12.16.423143

**Authors:** Mónica Durán, Carla Burballa, Gerard Cantero-Recasens, Cristian Butnaru, Vivek Malhotra, Gema Ariceta, Eduard Sarró, Anna Meseguer

## Abstract

Dent disease 1 (DD1) is a rare X-linked renal proximal tubulopathy characterized by low molecular weight proteinuria (LMWP) and variable degree of hypercalciuria, nephrocalcinosis and/or nephrolithiasis with progression to chronic kidney disease (CKD). Although loss-of-function mutations in the gene *CLCN5* encoding the electrogenic Cl^−^/H^+^ antiporter ClC-5, which impair endocytic uptake in proximal tubule cells, cause the disease, there is poor genotype-phenotype correlation and their contribution to proximal tubule dysfunction remains unclear. Here, in order to discover the mechanisms leading to proximal tubule dysfunction due to ClC-5 loss-of-function, we have generated and characterized new human cellular models of DD1 by silencing *CLCN5* and introducing the ClC-5 pathogenic mutants V523del, E527D and I524K into the human proximal tubule-derived cell line RPTEC/TERT1. Depletion of *CLCN5* or expression of mutant ClC-5 impairs albumin endocytosis, increases substrate adhesion and decreases collective migration, which correlates with a less differentiated epithelial phenotype. Interestingly, although all conditions compromised the endocytic capacity in a similar way, their impact on gene expression profiles was different. Our DNA microarray studies show that ClC-5 silencing or mutant re-introduction alter pathways related to nephron development, anion homeostasis, organic acid transport, extracellular matrix organization and cell migration, compared to control cells. Cells carrying the V523del ClC-5 mutation show the largest differences in gene expression vs WT cells, which is in agreement with the more aggressive clinical phenotype observed in some DD1 patients. Overall, this work emphasizes the use of human proximal tubule derived cell models to identify the molecular processes underlying ClC-5 deficiency.

## Introduction

Dent disease 1 (DD1; OMIM #300009) is a rare X-linked renal tubulopathy affecting about 330 families world-wide [1] and characterized by low molecular weight proteinuria (LMWP), and variable degree of hypercalciuria, nephrocalcinosis, calcium nephrolithiasis, and hypophosphatemic rickets [2,3]. DD1 progresses to renal failure between the 3rd and 5th decades of life in 30-80% of affected males, while female carriers are usually asymptomatic [4]. There is no current curative treatment for DD1 and patient’s care is supportive, focusing on the treatment of hypercalciuria and the prevention of nephrolithiasis [5]. DD1 is caused by loss-of-function mutations in the *CLCN5* gene encoding the electrogenic 2 Cl^−^/H^+^ antiporter ClC-5, which is abundantly expressed in the epithelia of kidney and intestine, though it is also expressed in brain, lung and, to a lesser extent, liver [6]. In the human kidney, ClC-5 is mainly expressed in proximal tubule cells (PTCs), where it is predominantly located in intracellular subapical endosomes and participates in endosomal acidification [2,7]. A small fraction of ClC-5 is also found on the plasma membrane of PTCs, where it is proposed to mediate plasma membrane chloride currents [7] or participate in the macromolecular complexes responsible for LMW protein and albumin endocytosis [8].

PTCs reabsorb approximately 65% of filtered load and most, if not all, of filtered LMW proteins mainly via receptor-mediated endocytosis [9]. The main actor in LMW protein reabsorption is the endocytic complex, which is comprised by the multiligand tandem receptors megalin and cubilin. Receptor-mediated endocytosis requires a continuous cycling of megalin and cubilin between the apical plasma membrane, where they specifically bind ultrafiltrated LMW proteins and other ligands, and the early endosome, where the receptors dissociate from their bound ligands [10]. This process requires vesicular acidification for dissociating the ligand-receptor complex, recycling of receptors to the apical membrane, and progression of ligands into the lysosomes. Endosomal acidification is achieved by ATP-driven transport of cytosolic H^+^ through the vacuolar H^+^-ATPase [11]. Inactivating mutations of *CLCN5* in Dent disease patients [12] as well as the deletion of *CLCN5* in knock-out (KO) mice [13,14] lead to severe LMWP due to a defective endocytic uptake in PTCs, which has been associated with the disappearance of megalin and cubilin at the brush border of PTCs. ClC-5 was initially postulated to provide a Cl^-^ shunt into the lumen of endosomes to dissipate V-ATPase-mediated H^+^ accumulation, thereby enabling efficient endosomal acidification [11,15]. However, mutations in ClC-5 causing Dent disease do not necessarily lead to a defective endosomal acidification [16], suggesting that the disease may result from an impaired exchange activity, namely uncoupling Cl^-^/H^+^ co-transport and altered Cl^-^ accumulation at early endosomes [17]. Thus, the precise molecular role of ClC-5 in endosomal physiology and endocytosis, as well as several aspects of its ion transport properties remain to be fully elucidated.

To date, a total of 266 pathogenic variants of *CLCN5* have been reported consisting of nonsense, missense, splice site, insertion and deletion mutations [1,18]. According to the latest reports, *CLCN5* mutations are grouped into three classes on the basis of functional data [16,18,19]: class 1 mutations result in defective protein processing and folding, thereby inducing retention of the mutant protein in the endoplasmic reticulum (ER), where they are early degraded by quality control systems; class 2 mutations impair protein processing and stability, leading to a functionally defective protein lacking electric currents; these mutants show reduced expression in the plasma membrane, but a normal distribution in the early endosomes; and class 3 mutations generate a protein that reaches the plasma membrane and early endosomes correctly, but shows reduced or abolished currents.

Yet, very little is known regarding how these mutations lead to specific disease manifestations. In this sense, the considerable intra-familial variability in disease severity and the lack of genotype-phenotype correlation suggest that unknown mechanisms might be involved in PTCs dysfunction leading to DD1 progression. In order to identify these ClC-5 mutation-associated pathways, we have silenced the *CLCN5* gene or introduced the ClC-5 mutations V523del (not classified), E527D (class 2) or I524K (class 1) in RPTEC/TERT1 cells. This cell line represents one of the most well-differentiated and stable proximal tubular cell line currently available, retaining sodium-dependent phosphate uptake and an intact functionality of the megalin/cubilin transport system [20,21]. Gene expression profiling and functional analysis in these cells revealed the biological processes related to proximal tubule dysfunction in DD1, likely explaining phenotype variability of the disease and the progression to renal failure.

## Results

### Expression and subcellular localization of ClC-5 mutants V523del, E527D and I524K in RPTEC/TERT1 cells

To explore the molecular mechanisms underlying PTCs dysfunction in DD1, first we have generated stable RPTEC/TERT1 cell lines silenced for *CLCN5* gene or carrying the pathogenic ClC-5 mutations V523del, E527D or I524K (described in DD1 patients [22–24]). We chose to study these mutations because, although their close location within the P helix of ClC-5, which is involved in dimer interface’s formation, and all three mutations resulting in loss of ClC-5 activity, they differentially affect ClC-5 subcellular localization and functionality [16,22,23]. I524K is a class I mutation and abolished currents have been related to its retention in the endoplasmic reticulum (ER) [16]. E527D is a type 2 ClC-5 mutant, and it lacks currents despite its normal presence in the endosome compartment and partially (30%) reaching the plasma membrane [16]. Expression of both E527D and I524K mutants in HEK293 cells also resulted in impaired endosomal acidification and altered protein stability [16]. V523del effects on subcellular localization and endosomal acidification have not yet been described.

A scheme summarizing the generation of the RPTEC/TERT1 DD1 cell model and the localization of shRNA sequences and mutations within ClC-5 is provided in figures S1 and S2. First, to fully characterize these cell lines, RNA was extracted from 10-day differentiated control, *CLCN5* knockdown (KD), rClC-5 WT, rClC-5 V523del, rClC-5 E527D and rClC-5 I524K carrying cells and endogenous *CLCN5* and exogenous ClC-5 (HA) levels monitored by real-time quantitative PCR (RT-qPCR). Our results showed that endogenous levels of ClC-5 were strongly reduced in all cell lines transduced with the shRNA against ClC-5 compared to control cells (6.2 %, 16.1%, 4.4%, 8.2% and 13% of control shRNA ClC-5 expression levels for *CLCN5* shRNA, rClC5 WT, rClC5 V523del, rClC5 E527D and rClC5 I524K, respectively) (Fig. 1A). Re-introduction of HA-tagged wild-type (rClC5 WT) or mutant (rClC5 V523del, rClC5 E527D and rClC5 I524K) ClC-5 in previously ClC-5 silenced cells restored ClC-5 mRNA levels above those of control cells (Ctrl shRNA) (Fig. 1B), although, rClC-5 V523del and rClC-5 E527D but not rClC5 I524K mRNA levels were lower than for the rClC-5 WT condition (33.8%, 68.7% and 87.9% of rClC-5 WT ClC-5 expression levels, respectively). At the protein level, ClC-5 was detected as a lower band running at 80-90 kDa and a higher diffuse band running as a smear at about 100 kDa, which was consistent with previous reports [25,26] (Fig. 1C). Loading equivalent amount of cell extract revealed that the protein levels of all the three ClC-5 mutants were strongly reduced in comparison with rClC-5 WT (Fig. 1C).

**Figure 1.**
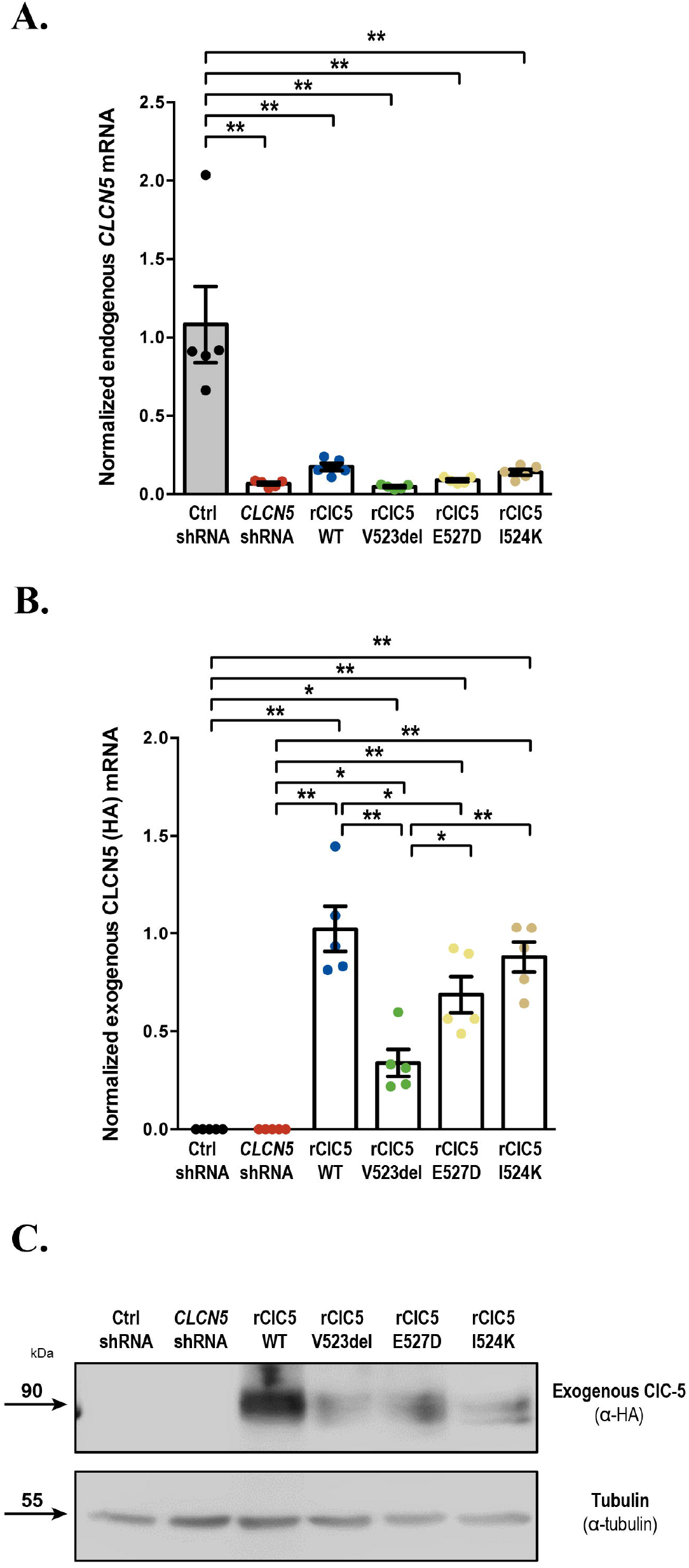
Expression levels of ClC-5 mutants V523del, E527D and I524K in RPTEC/TERT1 cells. DD1 cell models were generated by silencing *CLCN5* and re-introducing either wild-type (WT) or mutant (V523del, E527D and I524K) forms of ClC-5. To escape from RISC-mediated degradation, re-introduced ClC-5 forms incorporated silent mutations in the shRNA target sequence. (A) To validate that the *CLCN5* gene was silenced in all cell lines transduced with the *CLCN5* shRNA, mRNA levels of endogenous *CLCN5* were measured by RT-qPCR using specific probes targeting the intact shRNA target sequence. (B) The expression levels of re-introduced WT (rClC-5 WT) and mutant (rClC-5 V523del, rClC-5 E527D and rClC-5 I524K) ClC-5 were assessed by RT-qPCR using specific probes against the HA-tag, which is only present in the exogenous ClC-5. (C) Protein levels of re-introduced WT and mutant ClC-5 were analyzed by Western blot using an antibody against the HA-tag. Tubulin was used to ensure that equal amounts of total cell extract were loaded in each lane. For all experiments, Ctrl shRNA corresponds to cells transduced with both shRNA empty vector and re-expression empty vector. *, p < 0.05; **, p < 0.01.

We next analyzed the subcellular localization of WT and mutant ClC-5 in RPTEC/TERT1 cells by using immunostaining techniques. Our results show that rClC-5 WT localized at the plasma membrane (PM), early endosomes (EE) and endoplasmic reticulum (ER), as shown by co-localization with the specific subcellular compartment markers N-cadherin (PM), Rab-5 (EE) and KDEL (ER) (Fig. 2A, B and C). Quantification of the co-localization of ClC-5 forms with KDEL using the Manders’ overlap coefficient (MOC) demonstrated that rClC5 E527D (MOC = 0.29) and rClC-5 I524K (MOC = 0.39), but not rClC-5 V523del (MOC = 0.17) accumulated at the ER to a greater extent than rClC-5 WT (MOC = 0.10). Moreover, all three mutants showed a reduced co-localization with Rab5 (MOC rClC-5 WT = 0.29, rClC-5 V523del = 0.09, rClC5 E527D = 0.16 and rClC-5 I524K = 0.07) and N-cadherin (MOC rClC-5 WT = 0.33, rClC-5 V523del = 0.03, rClC5 E527D = 0.02 and rClC-5 I524K = 0.01) in comparison with rClC-5 WT, indicating that their presence in EE and PM was reduced. These results confirmed, in the case of E527D and I524K subcellular localizations, previous results in HEK-MSR cells [16].

**Figure 2.**
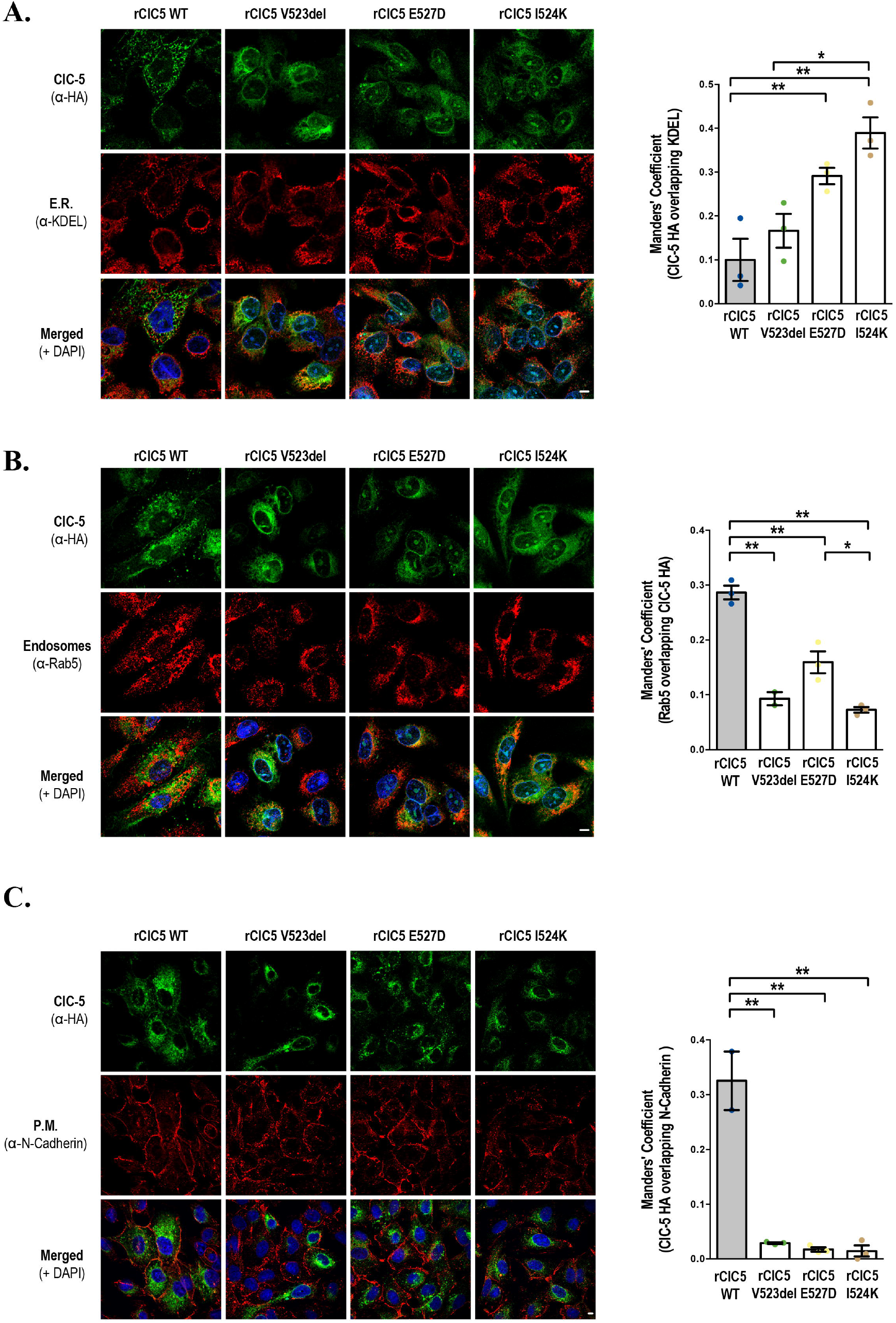
V523del, E527D and I524K mutations alter the subcellular localization of ClC-5 in RPTEC/TERT1 cells. Subcellular localization of WT (rClC-5 WT) and mutant (rClC-5 V523del, rClC-5 E527D and rClC-5 I524K) ClC-5 was analyzed in RPTEC/TERT1 cells seeded on glass coverslips by determining their co-localization with the endoplasmic reticulum (ER) marker KDEL (A), early endosomes (EE) marker Rab-5 (B) and plasma membrane (PM) marker N-cadherin (C), using the corresponding antibodies. Cell nuclei were stained with DAPI. Quantification of the co-localization was performed using the Manders’ overlap coefficient (MOC). *, p < 0.05; **, p < 0.01.

### I524K mutation, but not V523del or E527D presents an altered glycosylation pattern

It has been previously described that ClC-5 undergoes several post-translational modifications, including glycosylation [27]. Moreover, mutations on ClC-5 N-glycosylation sites trigger poli-ubiquitination and proteasomal degradation [26,27]. To gain more insight on whether the differences in protein levels and sub-cellular localization between rClC-5 WT and ClC-5 mutants could be related to impaired glycosylation processing, cell lysates from each of the conditions were treated with Endoglycosidase H (Endo H), which cleaves asparagine-linked mannose rich oligosaccharides, but not highly processed complex oligosaccharides, and Peptide:N-glycosidase F (PNGase F), which cleaves between the innermost GlcNAc and asparagine residues of high mannose, hybrid, and complex oligosaccharides. Our results show that rCLC-5 WT and all mutant forms of ClC-5 were sensitive to PNGase F digestion, confirming that all them were N-glycosylated (Fig. 3). On the other hand, only the lower migrating band of I524K mutant was sensible to EndoH digestion, as observed by a reduction in its molecular weight. These results suggest that this faster migrating band of the I524K mutant might correspond to a core-glycosylated EndoH-sensitive form of the protein, correlating with the higher degree of ER retention observed for this mutant.

**Figure 3.**
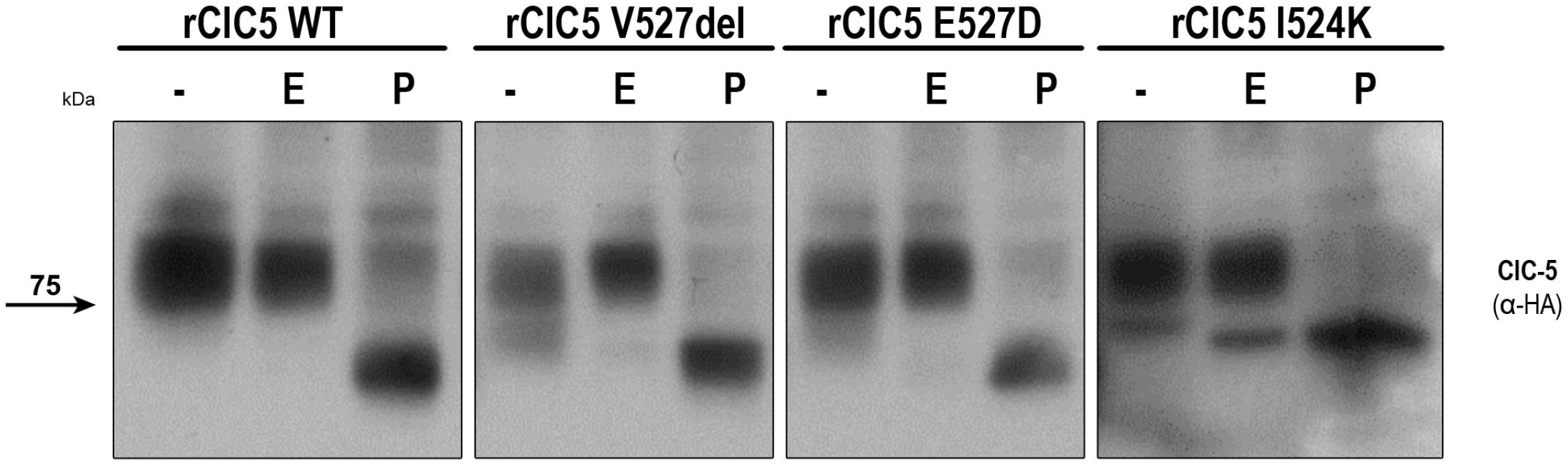
I524K ClC-5 mutant, but not V523del or E527D, presents an altered glycosylation pattern. To explore the effects of selected ClC-5 mutations on the glycosylation pattern of ClC-5, total cell lysates from RPTEC/TERT1 cells carrying each of the mutations were treated with Endoglycosidase H (E), which cleaves asparagine-linked mannose rich oligosaccharides, but not highly processed complex oligosaccharides, and Peptide:N-glycosidase F (PNGase F) (P), which cleaves between the innermost GlcNAc and asparagine residues of high mannose, hybrid, and complex oligosaccharides. After glycosidase reactions, samples were analyzed by Western Blot using an anti HA-tag antibody.

To investigate whether the expression of ClC-5 mutants, and more specifically, I524K, could be inducing the Unfolded Protein Response (UPR) and ER stress as a result of their accumulation in the ER, we checked the phosphorylation of the ER stress marker PERK [28] (Fig. S2A) and the cleavage of XBP-1 mRNA [28] (Fig. S2B) in cells expressing WT or mutant ClC-5. Our results show that neither expression of rCLC-5 WT nor any of the ClC-5 mutants studied induced detectable levels of ER stress.

### ClC-5 silencing or re-introduction of V523del, E527D and I524K ClC-5 mutants impairs Albumin endocytosis

To determine the effect of ClC-5 silencing and the selected ClC-5 mutations on the endocytic capacity of RPTEC/TERT1 cells, we analyzed Alexa Fluor 488-labelled albumin uptake. Detection of labeled albumin within the cell boundaries, both in orthogonal views and single planes, of control cells demonstrated that the endocytic machinery functions properly in the RPTEC/TERT1 cell line (Fig. 4A). ClC-5 silencing strongly reduced the uptake of labelled albumin, whereas the endocytic capacity was re-established when the wild-type ClC-5 was re-introduced in the silenced cells (number of albumin particles/cell Ctrl shRNA = 7.23, *CLCN-5* shRNA = 2.27 and rClC-5 WT = 6.57) (Fig. 4A and B). By contrast, re-introduction of neither V523del nor E527D nor I524K ClC-5 mutants was unable to restore this activity (number of albumin particles/cell rClC-5 V523del = 3.15, rClC-5 E527D = 3.7 and rClC-5 I524K = 2.58), what indicates that these residues are essential for ClC-5-mediated endocytosis (Fig. 4A and B). In addition, the volume of particles (which is related to the amount of endocytosed albumin) was also reduced by depleting *CLCN5* (volume of particles Ctrl shRNA = 0.19 and *CLCN-5* shRNA = 0.09) or expression of loss-of-function ClC-5 mutants (volume of particles rClC-5 WT = 0.24, rClC-5 V523del = 0.11, rClC-5 E527D = 0.13 and rClC-5 I524K = 0.08) (Fig. 4C).

**Figure 4.**
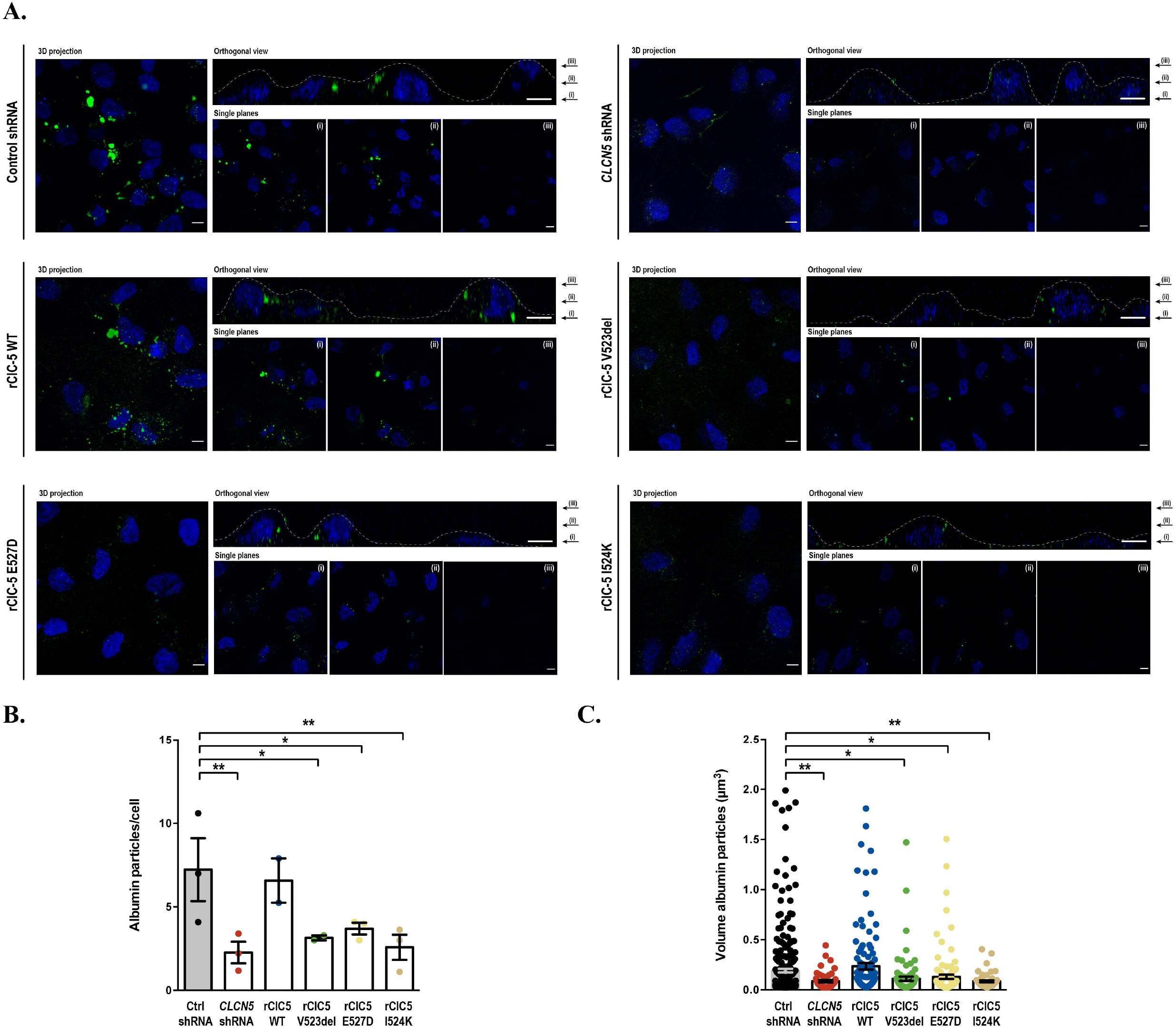
ClC-5 silencing or V523del, E527D and I524K ClC-5 mutations impair Albumin endocytosis. To determine the effects of ClC-5 silencing and the selected ClC-5 mutations on the endocytic capacity of RPTEC/TERT1 cells, we analyzed Alexa Fluor 488-labelled albumin uptake. (A) Cells were seeded on glass coverslips and incubated with 50 μg/mL Alexa Fluor 488-conjugated Albumin for 60 min. After extensive washing, endocytosed albumin was detected within the cells. In the orthogonal view, dotted lines across the images demarcates the top surface of the cell. Quantification of albumin uptake was performed by measuring the number of albumin particles/cell (B) and the volume of albumin particles (C), which is related to the amount of endocytosed albumin. *, p < 0.05; **, p < 0.01.

### ClC-5 silencing or re-introduction of ClC-5 mutations V523del, E527D and I524K alter the global gene expression profile of RPTEC/TERT1 cells

In order to discover potential mechanisms involved in the proximal tubule dysfunction secondary to the loss of ClC-5, we analyzed the gene expression profile of DD1 cell models using DNA microarrays. To make the data comparable, as well as to remove technical biases, microarray data were first normalized and batch effect corrected. The Principal Component Analysis (PCA) obtained after applying these corrections is shown in figure 5A, where it can be observed that samples were mainly grouped by condition. In addition, and to validate the reliability of the results obtained from the DNA microarray, the expression levels of EMX2, PTPRD, STEAP1, ZPLD1, CDH1, NR1H4 and EHF genes were analyzed by qRT-PCR (Fig. S3). Validation genes were selected among those that meet the following requirements: i) their expression was altered by some of the mutations compared to the WT condition and, ii) its expression was also modified by the silencing of *CLCN5* and totally or partially restored by ClC-5 WT re-introduction. Our results show that all the validation genes presented an expression pattern similar to that in the DNA microarray (Fig. S3), thereby confirming the trustworthiness of the microarray data.

**Figure 5.**
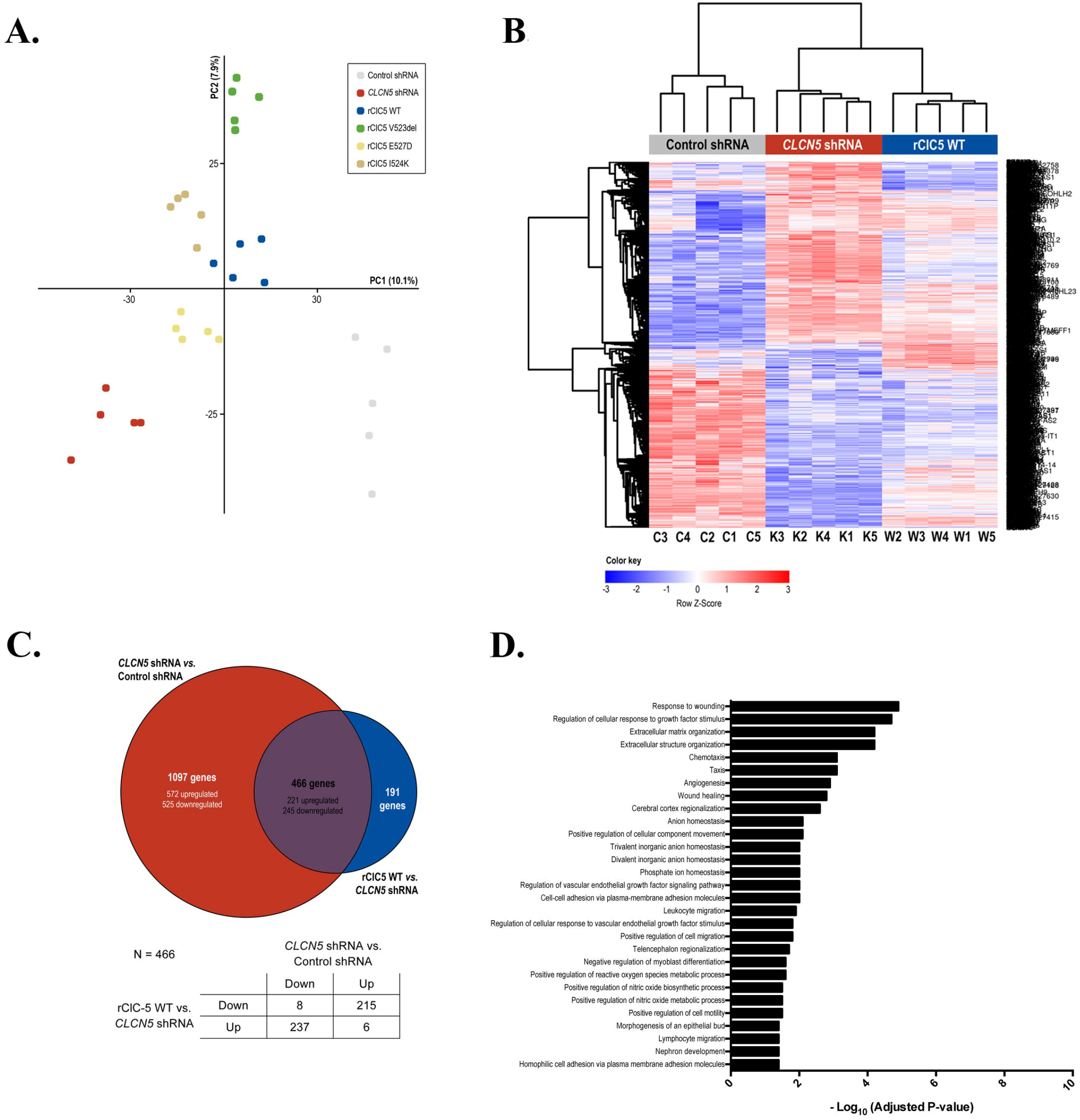
ClC-5 silencing alter the global gene expression profile of RPTEC/TERT1 cells. To identify the potential mechanisms involved in the proximal tubule dysfunction secondary to the loss of ClC-5, we analyzed the gene expression profile of RTPEC/TERT1 cells carrying *CLCN5* silencing or rClC-5 WT using DNA microarrays. (A) Principal Component Analysis (PCA) obtained after normalization and batch effect corrections of the DNA microarray data. (B) Heatmap graphically illustrating the differences in the gene expression profile of ClC-5-silenced cells *(CLCN5* shRNA) in comparison to control cells (control shRNA) and ClC-5-silenced cells where WT ClC-5 was re-introduced (rClC-5 WT). (C) Venn diagrams depicting the genes commonly regulated by the effect of *CLCN5* silencing and rClC-5 WT re-introduction. The table indicates the number of common genes that are up- or down-regulated in each comparison. (D) Analysis of over-represented gene ontology (GO) terms biological processes (BP) related to the genes commonly up- or down-regulated by the effect of *CLCN5* silencing and rClC-5 WT re-introduction, using the GProfiler server.

Figure S4 shows the number of genes in the DNA microarray whose expression was altered within a range of logFC for an adjusted p value lower than 0.05 in each of the comparisons that we have performed in this work. For comparative analysis, genes were defined as differentially expressed genes (DEGs) if they presented an adjusted p value lower than 0.05 and a log2 Fold Change (logFC) higher or equal to 0.5 in any of the comparisons studied. Top up- and down-regulated DEGs in each of the comparisons are shown in Supplementary Tables S1-8.

The Heatmap in Figure 5B graphically illustrates the differences in the gene expression profile of ClC-5-silenced cells in comparison to control cells and ClC-5-silenced cells with WT ClC-5 re-introduced. These results show that ClC-5 silencing elicited a marked effect on the gene expression profile of RPTEC/TERT1 cells, and that this effect was partially reversed by re-introducing the rClC-5 WT (Fig. 5B). Notably, from the 1563 genes altered by ClC-5 silencing, up to 452 (from a total of 466 commonly regulated genes) were regulated in the opposite direction by ClC-5 re-introduction (Fig. 5C). Among these, 237 (52%) genes were down-regulated and 215 (48%) were up-regulated by the effect of ClC-5 silencing. Only the genes that were commonly regulated by ClC-5 silencing and rClC-5 WT re-introduction (452 genes) were considered for subsequent analysis. To study in which biological processes (BP) were involved these genes, we performed an analysis of over-represented gene ontology (GO) terms. A list of the biological process GO terms significantly enriched is shown in Figure 5D. We identified GO terms related to (intersection size/term size): response to wounding (36/372), matrix organization (27/397), anion homeostasis (5/65), cell adhesion (16/275), cell migration (29/569), positive regulation of reactive oxygen species (ROS) (12/107) and nephron development (13/146), among others. In anion homeostasis significantly changed transcripts were SLC34A2, NR1H4, TFAP2B, SFRP4 and SLC7A11, and in nephron development were ADAMTS16, TFAP2B, NOG, FMN1, LGR4, DLL1, COL4A4, KIF26B, SULF1, NID1, TACSTD2, PROM1 and BMP4.

In order to explore the effects of the selected ClC-5 mutations on the transcriptome of RPTEC/TERT1 cells, gene expression profiles of V523del, E527D and I524K mutants were compared to that of WT ClC-5. As shown in the heatmap in figure 6A, V523del and I524K were the conditions that exhibited the greatest and the smallest differences in the gene expression profile, respectively, when compared to WT ClC-5. Moreover, only 5 genes (CHCHD7, PREX1, PLAG1, LIN54 and ZRANB3) were commonly affected by all three mutations (Fig. 6B). Re-introduction of V523del mutant altered the expression of 831 genes compared to ClC-5 WT cells, from which 231 were down-regulated and 600 were up-regulated. Roughly 20% (47/231) of the down- and 5% (27/600) of the up-regulated genes by Val523del expression were also found down- or up-regulated, respectively, in ClC-5-silenced cells, and might represent genes whose expression levels cannot be restored by V523del mutant to the levels achieved by WT ClC-5. On the other hand, genes altered by V523del that were not affected in ClC-5 silenced cells could be indicating a gain of functionality of this mutant with regard to ClC-5 silencing. An analysis of the significantly enriched biological processes (GO terms) showed that V523del altered the expression of genes mainly related to DNA replication, but also related to carboxylic acid and anion transport and renal system development, among others (Fig. 6C). ClC-5 mutant E527D altered the expression of 510 genes (204 down-regulated and 306 up-regulated). Among them, 24% of the down- (49/204) and 16% (48/306) of the up-regulated genes were also found down- or up-regulated, respectively, by effect of ClC-5 silencing. Genes whose expression was significantly altered by E527D in comparison to WT ClC-5 were specially enriched in processes related to tissue remodeling and morphogenesis, but also in other processes including (intersection size/term size): sialic acid transport (3/7), sodium ion transmembrane transport (8/152) and also renal system development (19/315), for example (Fig. 6D). Finally, I524K ClC-5 mutation altered the expression of only 32 genes compared to ClC-5 WT, with 6 of them down-regulated and 26 up-regulated. This low number of genes did not allow to obtain any significantly enriched GO term in the over-representation analysis.

**Figure 6.**
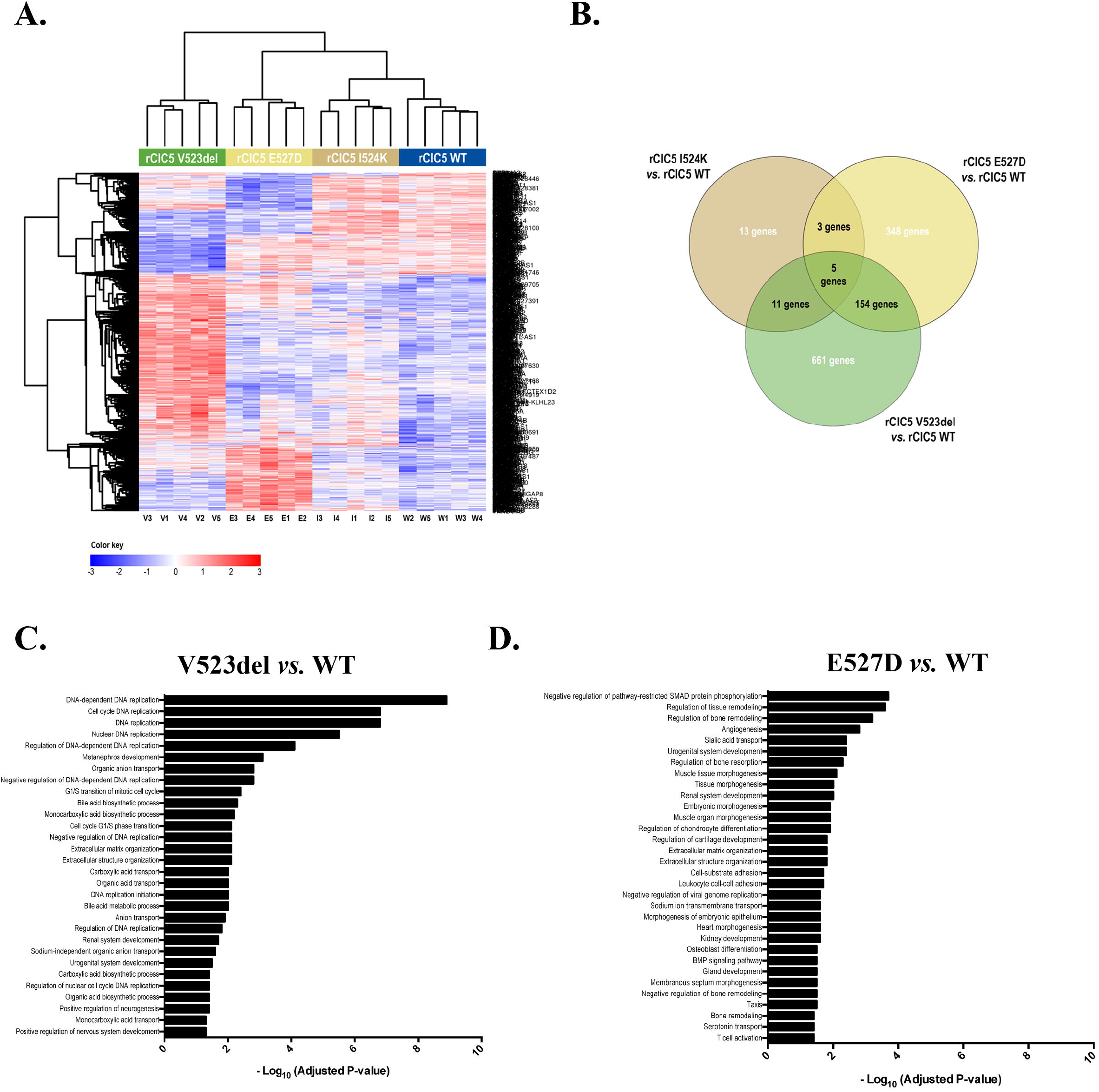
ClC-5 mutations V523del, E527D and I524K exert different effects on RPTEC/TERT1 gene expression profile. In order to explore the effects of the selected ClC-5 mutations on the transcriptome of RPTEC/TERT1 cells, gene expression profiles of V523del, E527D and I524K mutants were compared to that of WT ClC-5. (A) Heatmap showing that V523del and I524K were the conditions that exhibited the greatest and the smallest differences in the gene expression profile, respectively, when compared to WT ClC-5. (B) Venn diagrams depicting the number of commonly regulated genes by ClC-5 mutations. Only 5 genes were shown to be commonly affected by all three mutations. Analysis of the significantly enriched biological processes associated with the genes up- or down-regulated by V523del (C) or E527D (D) mutations in comparison to ClC-5 WT.

### ClC-5 silencing and mutations V523del, E527D and I524K impair cell-to-substrate adhesion and collective cell migration

We next explored whether the changes observed in the gene expression profiles by ClC-5-silencing or loss-of-function ClC-5 mutations correlated with changes in the epithelial characteristics (e.g. substrate adhesion or cell migration). For this purpose, we analyzed epithelial markers’ levels (e.g. CDH1, occludin, and Keratin-7 and −18), cell proliferation, substrate adhesion and collective cell migration. Results in Figure 7A show that ClC-5 silencing strongly reduced E-Cadherin and Keratin-7 levels, while it had no effect on occludin and Keratin-18 levels. Re-introduction of rClC-5 WT, but not ClC-5 mutants V523del, E527D or I524K, totally rescued E-cadherin and keratin-7 expression (Fig 7A). Our results also show that, in comparison to control cells, ClC-5 silencing increased cell-to-substrate adhesion (Fig. 7B), reduced collective cell migration (Fig. 7C) and, although not in a statistically significant manner, increased cell proliferation (Fig. 7D). In a similar way, all three ClC-5 mutants increased cell-to-substrate adhesion (Fig. 7E) and reduced collective cell migration (Fig. 7F) when compared to the respective control rClC-5 WT. By contrast, no apparent differences were observed in the proliferation rates of ClC-5 mutant cell lines compared to cells carrying the WT protein (Fig. 7G). Taken together, these results suggest that both ClC-5 silencing and expression of ClC-5 mutants could lead cells into a dedifferentiated state.

**Figure 7.**
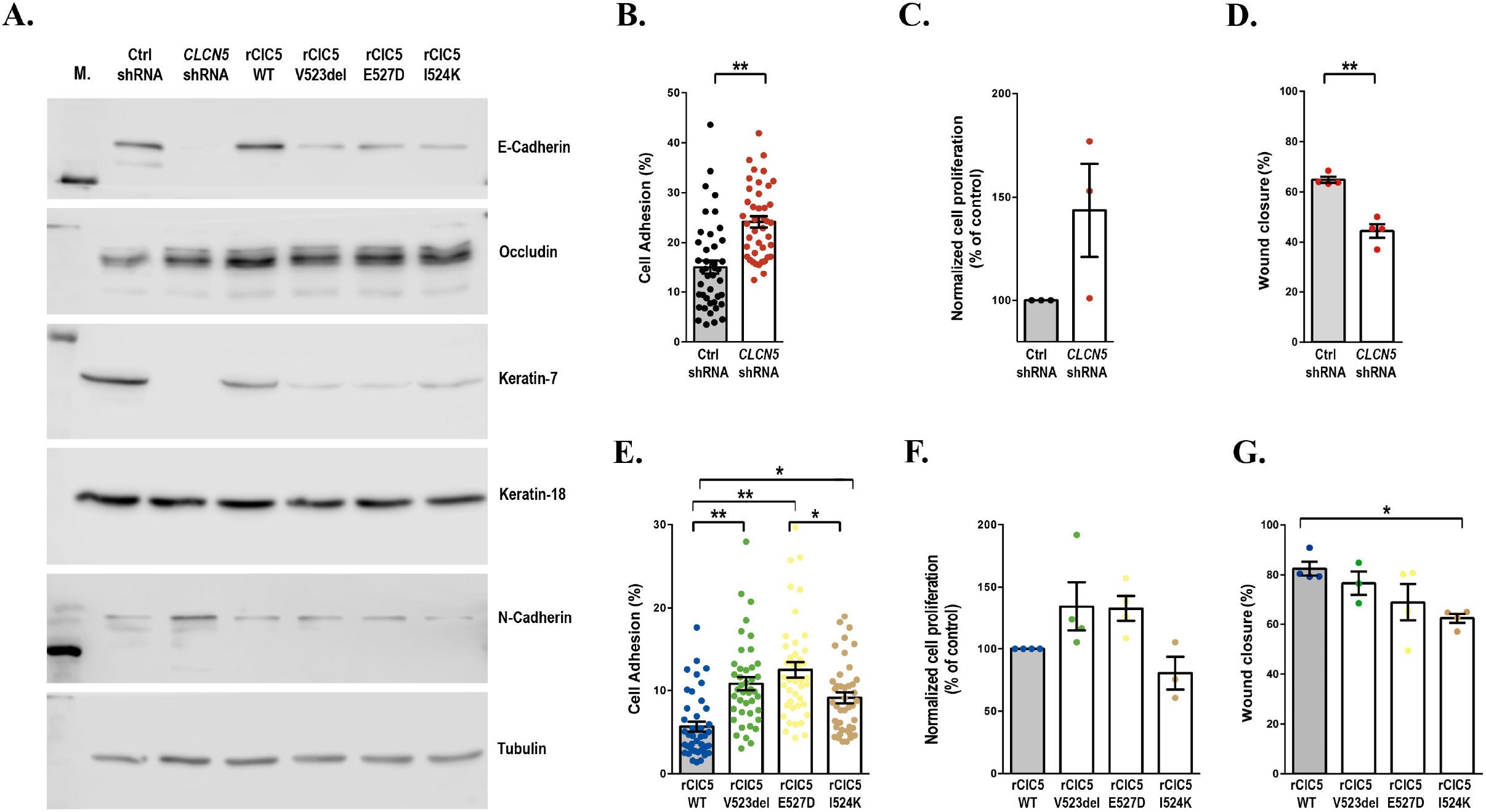
ClC-5 silencing and mutations V523del, E527D and I524K impair cell-to-substrate adhesion and collective cell migration. To explore whether the changes observed in the gene expression profiles by ClC-5-silencing or loss-of-function ClC-5 mutations correlated with changes in the epithelial characteristics, we analyzed substrate adhesion, proliferation and collective cell migration in RTEC/TERT1 cells. (A) Epithelial markers CDH1, occludin, and Keratin-7 and −18 were analyzed by Western Blot. (B and E) Cell-to-substrate adhesion as assessed by the ability of RPTEC/TET1 cells to bind to tissue culture substrate (B). (C and F) Cell proliferation was measured by staining cells with CFSE and quantifying the fluorescence of the cells at the onset of the experiment and 4 days later. (D and G) collective cell migration, which depends on the integrity of cell-cell contacts, was determined with the Wound healing assay as indicated in methods. *, p < 0.05; **, p < 0.01.

## Discussion

Besides reports describing the effects of *CLCN5* mutations on chloride currents and endosomal acidification, there is a paucity of studies addressing the impact of these mutations on the phenotype and expression profile of PTCs. Consequently, little is known about the potential mechanisms involved in the proximal tubule dysfunction secondary to the loss of ClC-5. Moreover, evidences indicate that CLC-5 mutations causing defective proximal tubular endocytosis and endosomal trafficking may not necessarily impair endosomal acidification, suggesting that these processes may not be coupled ant that tubular dysfunction in DD1 may not result from reduced endosomal acidification [16,29]. In this sense, one of the most striking conclusions of this study is that despite the lack of ClC-5 or the presence of ClC-5 mutations V523del, E527D and I524K compromised the endocytic capacity of RPTEC/TERT1 in a similar way, they do not exert equivalent effects on the gene expression profile of these cells, and only five genes were commonly modulated by these mutations. This suggests that mutant forms of ClC-5 may alter different cellular processes apart from the endocytic pathway. Accordingly, in this work we aimed to discover the mechanisms that underlie tubule dysfunction in DD1 by characterizing the phenotypical consequences of ClC-5 mutations on PTCs.

For this purpose, we have generated novel Dent disease cell models by stably transfecting *CLCN5* shRNA and pathogenic ClC-5 mutations into the RPTEC/TERT1 cell line, which maintains many differentiation hallmarks [20]. The use of cell lines takes advantage of an uniform genetic background thus avoiding compensatory effects due to currently unknown polymorphisms or mutations in other genes. In addition, use of cell models is of special interest in Dent disease, where renal biopsy are not routinely indicated since i) laboratory findings and genetic testing can be sufficient for diagnosis and ii) most DD1 patients are children and it’s not worth applying an invasive procedure if it cannot provide relevant information. In these cell models, we have characterized ClC-5 mutant proteins and have identified genes and biological processes specifically regulated by ClC-5 silencing or re-introduction of mutants forms of ClC-5. In this regard, we found that RPTEC/TERT1 cells lacking ClC-5 showed a marked loss of epithelial markers, an increase in cell to substrate adhesion, reduced collective cell migration and a trend suggesting an increase in cell proliferation, all of them characteristics of epithelial dedifferentiation. This correlated with an altered expression of genes related to reactive oxygen species, cell-cell adhesion, cell migration, extracellular matrix organization or cell motility among others. In the same direction, Gally et al [30] found that PTCs taken from ClC-5 KO mice had increased expression of proliferation markers and oxidative scavengers, suggesting that PT dysfunction in ClC-5 KO mice was associated with oxidative stress, dedifferentiation and increased cell proliferation. Moreover, the urinary proteome of patients with Dent disease has been shown to be enriched with proteins actively participating in interstitial matrix remodeling [31]. Thereby, our results are aligned with the hypothesis raised by Devuyst and Luciani [17] explaining the potential mechanism by which the loss of ClC-5 may cause proximal tubule dysfunction. According to the authors, and in addition to the impaired trafficking and recycling of apical receptors and the defective receptor-mediated endocytosis, loss of ClC-5 would be also associated with altered lysosomal function. This might compromise the lysosomal mediated-degradation and clearance of autophagosomes containing ubiquitinated proteins and dysfunctional mitochondria, leading to excessive production of reactive oxygen species (ROS). The increase in ROS might alter the integrity of the junctional complex proteins, releasing transcription factors, which will translocate to the nucleus and promote proliferation. It is noteworthy that we did not detect a significant enrichment of GO terms related to phagocytosis in neither ClC-5 silenced cells nor cells carrying V523del, E527D nor I524K ClC-5 mutations, thereby suggesting the existence of other molecular mechanisms converging on PTCs dedifferentiation and dysfunction. In this sense, and in addition to the abovementioned GO terms, ClC-5 silencing also altered the expression of genes related to biological processes such as anion homeostasis, chemotaxis or response to growth factor. Moreover, terms such as morphogenesis of an epithelial bud and nephron development point at a role of ClC-5 in kidney development. It is worth mentioning that GO terms found in ClC-5-silenced RPTEC/TERT1 cells including organ development, ion transport, response to external stimulus, response to wounding, regulation of cell differentiation, chemotaxis and taxis were also found in the gene microarray analysis from proximal S1 and S2 tubules of ClC-5 KO mouse kidneys mice of the Guggino group [32], indicating that our cell model mimics the PTCs of the ClC-5 KO mouse. By contrast, we did not found terms related to lipid metabolism, which was the class with the greatest number of changes in gene transcript level in the ClC-5 KO mice [32]. That result was surprising because overall changes in lipids have not been reported in Dent disease. When we analyzed the list of genes altered by ClC-5 silencing, we found that among the most down-regulated genes there were genes that could be relevant in relation to DD1. Such an examples are SLPI (Secretory Leukocyte Peptidase Inhibitor; logFC −5.01), which has been related to PTCs regeneration [33], MUC1 (Mucin1; logFC −4.72), whose mutation causes a rare form of tubulointerstitial fibrosis [34], SLC34A2 (Sodium-dependent phosphate transport protein 2B; logFC −4.01), which may contribute to the diminution in the uptake of both sodium and phosphate in the proximal tubules in Dent disease patients, the Rab GTPase RAB27B (logFC −1.50), which is involved in exosome secretion, COL4A4 (Collagen Type IV Alpha 4 Chain; logFC −0.92), which is mutated in patients with Alport syndrome, the kidney-Specific Cadherin CDH16 (logFC −2.25), which is involved in cell-cell adhesions or KLF4 (Kruppel Like Factor 4; logFC −0.99) which has been identified as a renal linage master regulatory transcription factor [35]. Taken together, these results suggest that lack of ClC-5 widely affects the phenotype of RPEC/TERT1 cells, but it remains to be known whether lack of ClC-5 impacts on these processes through its effect on endosomal acidification, altered chloride transport, protein endocytosis, or its participation in macromolecular complexes. For instance, endocytic trafficking contributes to cell adhesion and migration in different ways [36]. First, internalization of chemokines by scavenger receptors is essential for sensing the chemotactic gradients, whereas endocytosis and subsequent recycling of chemokine receptors is key for sustaining the responsiveness of migrating cells. Second, endosomal pathways modulate adhesion by delivering integrins to their site of action and supplying factors for focal adhesion disassembly. Finally, endosomal transport also contributes to cell migration by delivering membrane type 1 matrix metalloprotease to the leading edge facilitating proteolysis-dependent chemotaxis.

One of the most striking results of the present work is the reduced number of biological processes commonly altered by ClC-5 silencing and re-introduction of ClC-5 mutations, and between each of the mutations, even though all conditions impaired albumin endocytosis and cell differentiation. Moreover, that’s despite their close location in ClC-5’s helix P and the fact that amino acids V523 and E527 are highly conserved residues present in all known ClCs [16,22,23]. Thus, only the GO terms “extracellular matrix organization” and “extracellular structure organization” were commonly found in the ClC-5 silencing and V523del and E527D conditions, while only the GO terms “urogenital system development” and “renal system development “ were common in V523del and E527D, although the term “nephron development” appeared in the ClC-5 silencing condition. Moreover, only 5 genes were commonly altered by all three mutations (CHCHD7, PREX1, PLAG1, LIN54 and ZRANB3). Accordingly, this lack of a functional equivalence between the absence of ClC-5 or the presence of ClC-5 mutants in relation to the biological processes point to a gain of functionality of the mutated forms of ClC-5.

Interestingly, V523del was the condition, among the different mutants studied, with the largest differences in gene profile when compared to the wild-type form. This could explain, in part, that V523del ClC-5 mutation has been found in a pediatric patient with a severe clinical phenotype [37], although the lack of more individuals carrying the same mutation makes impossible to establish such a correlation. Genes altered by V523del ClC-5 mutation were mainly involved in cell cycle and proliferation, which are processes that have been linked to a dedifferentiation state, but also in carboxylic acid and anion transport, and renal system development biological processes. Unexpectedly, cells carrying V523del ClC-5 only showed a small non-significant increase in cell proliferation compared to rClC-5 WT cells, thereby suggesting that the V523del-modulated genes included in proliferation GO terms could indeed be mediating dedifferentiation. On the other hand, we found altered an elevated number of genes from the solute carrier (SLC) group of membrane transport proteins. SLC transporters show high expression levels in metabolically active organs such as the kidney, liver or brain [38], and the kidney has been identified as one of the target organs for most high expression of SLCs-mediated diseases [39]. For instance, we found up-regulated the type I sodium-dependent phosphate transporters SLC17A1 (NPT1) and SLC17A3 (NPT4), which are the two most up-regulated genes in this condition, and also SLC27A2 (Fatty Acid Transporter FATP2), SLC16A4 (Monocarboxylate Transporter 4 MCT4) and SLC4A4 (Sodium Bicarbonate Cotransporter NBC1), all of them being involved in renal diseases [39]. To cite some, SLC17A1 and SLC4A4 mutations cause Fanconi Syndrome. By contrast, neither ClC-5 silenced cells nor any mutant condition showed an altered expression of the sodium-bile acid cotransporter SLC10A2, which was one of the gene transcripts most increased in transcript number (17 fold) in the *CLCN5* knockout mice proximal tubules of the Guggino group [32]. However, we found that V523del cells had a significant enrichment of the biological process “bile acid synthesis and transport”, and genes contained in this GO term, such as the bile acid receptor NR1H4, and the nuclear receptor NR1D1 or the Very Long-Chain Acyl-CoA Synthetase SLC27A2, were also among the most up-regulated genes in V523del cells. Thus, the fact that V523del ClC-5 up-regulates so many genes codifying for apical and basolateral membrane co-transporters may suggest the existence of compensatory pathways to overcome the defective receptor-mediated endocytosis caused by ClC-5 loss-of-function. Besides that, and as mentioned above, renal development was another biological process altered in V523del cells. Amid the genes belonging to this biological process, we found highly up-regulated (logFC = 2.31) the transcription factor HES1 (hairy and enhancer of split-1), since it has been previously identified as a renal linage master regulatory transcription factor, playing an important role in the Notch signaling pathway [35]. As for V523del down-regulated genes, it is remarkable to note that much of the most down-regulated genes, such as CDH1, MFAP5 or LUM, participate in the extracellular matrix organization. This effect could in part explain the reduced collective cell migration rates of the cells carrying the V523del mutation. Finally, it is also worth mentioning that cells carrying V523del mutation downregulate SLC3A1 gene, which codifies for the amino acid transporter ATR1 and is found mutated in cystinuria patients [39].

As mentioned earlier, E527 is one of the most conserved amino acids and is present in all the known ClCs, including those from plants, yeast, Escherichia coli, cyanobacteria, fish and mammals [16,23]. In addition, it has been previously described that the E527D ClC-5 mutant has a dominant negative effect on endosomal acidification [16], and mutation of the corresponding residue in ClC-0 results in a reversion of voltage dependence, i.e currents were activated by hyperpolarization instead of depolarization [40]. It is striking that, in our cell model, a large part of the biological processes altered after introduction of the E527D mutation were related to tissue remodeling, morphogenesis, differentiation and development. Interestingly, the biological processes “Organ development” and “organ morphogenesis” were the 2^nd^ and 8^th^ GO terms, respectively, with the greatest number of significantly changed transcripts in the ClC-5 KO mice of the Guggino group [32]. Other biological processes, such as sialic acid transport, sodium ion transmembrane transport and, cell substrate adhesion, BMP signaling pathway or T cell activation appeared altered in RPTEC/TERT1 cells carrying E527D ClC-5. In this sense, genes related to T cell activation, such as genes of the human leukocyte antigen (HLA) system (HLA-DRB1, logFC 1.79; HLA-DMA, logFC 1.55; and HLA-DPA1, logFC 1.44) or the lymphocyte cytosolic protein 1 (LCP-1, logFC −1.46) and Thy-1 cell surface antigen (THY1, logFC −1.44) are among of the most up or down regulated genes by E527D re-introduction. These results are consistent with microarray data obtained from intestine of ClC-5 KO mice showing altered expression of genes implicated in the immune system [41], and with the proposed role for ClC-5 in the immunopathogenesis of ulcerative colitis [42]. In addition, we also found biological pathways related to bone remodeling. Since bone homeostasis is tightly connected to phosphate metabolism, an enrichment of such biological processes could be indeed reflecting an altered phosphate regulation in PTCs. Moreover, alteration of these processes in PTCs could be related to the increased bone turnover previously described in the ClC-5 KO mouse model of Dent’s disease, likely explaining the propension to altered bone homeostasis in young Dent’s patients [43].

Curiously, introduction of the ClC-5 mutant I524K in RPTEC/TERT1 cells only altered the expression of a reduced number of genes, yielding, among the different mutants studied, the gene expression profile that more closely resembled that of the rClC-5 WT condition. These results were unexpected considering that i) the I524K mutant presented the highest degree of ER localization among the different ClC-5 mutants studied, and ii) the lack of a correspondence between I524K mRNA and protein levels might indicate that this mutant exhibits reduced protein stability or impaired post-translational processing. The increased ER retention of I524K, however, does not translate to an induction of the UPR, suggesting that I524K would be rapidly targeted to proteasomal degradation without displaying and ER stress gene signature. Proteasomal degradation, in turn, could explain the reduced levels of I524K protein that we detected by Western blot. A possible explanation for the reduced phenotypic effects of I524K would be that, although most of the proteins would be retained in the ER, a small but sufficient amount of the I524K proteins would manage to escape from the ER and reach its functional localization. This hypothesis would imply that I524K is, beyond its retention in the ER, a functional protein able to produce chloride currents. The reduced number of genes modulated by I524K did not allow to find statistically enriched GO terms. However, an analysis of the DEGs in I524K cells shed some light about the potential processes altered by this mutation. In this sense, the top down-regulated genes, CDH1 (E-cadherin) and KRT7 (keratin-7), correspond to well-established epithelial markers. This expression profile is aligned with the reduced cell to substrate adhesion and collective cell migration observed in rClC-5 I524K cells. Moreover, the third most down-regulated gene, CATSPER1, corresponds to a voltage-gated calcium channel, and the fifth, SLC38A8, to a putative sodium-dependent amino-acid/proton antiporter. On the other hand, PREX1, a guanine nucleotide exchange factor for RAC1, and the ferroxidase enzyme Ceruloplasmin (CP) were the most up-regulated genes by I524K. PREX1, which is one of the few gens altered by all the three ClC-5 mutations studied, has been identified as an important factor in tumor cell invasion and metastasis in a number of cancer models [44]. As for CP, it has been described that it plays an important role in cellular iron homeostasis and could protect kidney against a damage from iron excess [45]. Interestingly, CP was also found up-regulated in KO mice of the Guggino group, and the molecular function “iron ion binding” appeared in sixth position in the GO miner analysis of DEGs [32].

In conclusion, in this work we have generated new cell models of Dent disease that accurately reproduce ClC-5 defects and we have demonstrated that, besides the established critical function of ClC-5 in endocytosis, there are other mutation-associated pathways that could be relevant for the etiopathogenesis of DD1, likely explaining the phenotypic variability of DD1 patients. In this sense, we found that biological processes related to kidney development, anion homeostasis, organic acid transport, extracellular matrix organization and cell migration, were among the pathways that more likely could explain the pathophysiology of Dent disease 1.

## Declarations

Compliance with Ethical Standards.

## Acknowledgments

We thank the patient advocacy group Asdent (Asociación de la Enfermedad de Dent, https://www.asdent.es) for its continuous support. We thank all members of the Meseguer Lab for valuable discussions. Cell cytometry was carried out at the High Technology Unit at Vall d’Hebron Research Institute (VHIR). Fluorescence microscopy was performed at the Advanced Light Microscopy Unit at the CRG, Barcelona; and at the High Technology Unit at Vall d’Hebron Research Institue (VHIR). V.Malhotra is an Institució Catalana de Recerca i Estudis Avançats professor at the Centre for Genomic Regulation. This work reflects only the authors’ views, and the EU Community is not liable for any use that may be made of the information contained therein.

## Methods

### Cell culture

Renal proximal tubule epithelial cells RPTEC/TERT1 were obtained from the American Type Culture Collection (ATCC^®^; #CRL-4031). RPTEC/TERT1 were cultured in Dulbecco’s Modified Eagle Medium: Nutrient Mixture F-12 (1:1, v/v) (Thermo Fisher Scientific, #31331093) supplemented with 20 mM HEPES (Gibco, #15630-080), 60 nM sodium selenite (Sigma Aldrich, #S9133), 5 μg/ml transferring (Sigma Aldrich, #T1428), 50 nM dexamethasone (Sigma Aldrich, #D8893), 100 U/ml penicillin and 100 μg/ml streptomycin (Gibco, #15240-062), 2% fetal bovine serum (Gibco, #10270), 5 μg/ml insulin (Sigma Aldrich, #I9278), 10 ng/ml epidermal growth factor (Sigma Aldrich, #E4127)) and 3 nM triiodothyronine (Sigma Aldrich, #T5516). Cultures were maintained at 37 °C in a 5% CO_2_ atmosphere. Unless otherwise indicated, cell were cultured for 10 days to allow cell differentiation.

### Gene silencing

For *CLCN5* silencing, the MISSION^®^ TRC shRNA transfer vector containing the ClC-5 shRNA target sequence CACCGAGAGATTACCAATAA (Sigma-Aldrich, #TRCN0000043904) was co-transfected with the third generation vectors VSVG, RTR2 and PKGPIR, which provide the envelope, packaging and reverse-expressing proteins, respectively, into HEK-293 cells. Supernatants containing viral particles were then harvested, supplemented with 10% FBS, 1% non-essential amino acids and 8 μg/mL polybrene (Sigma-Aldrich, #TR-1003) and added to RPTEC/TERT1 cells, followed by antibiotic-mediated selection (8 μg/mL puromycin, Invivogen, #ant-pr).

### Vectors and Site-directed mutagenesis

For shRNA rescue experiments, wild-type human *CLCN5* was cloned into pDONR vectors (pDONR™221, Invitrogen, #12536-017) using the Gateway cloning system (Invitrogen). In order to escape from degradation by the RISC complex, silent mutations (c.[99C>T; 100C>A; 102A>G; 105G>A; 108T>C; 111C>A; 114T>C]) were introduced in the shRNA targeting sequence of human *CLCN5.* Over the shRNA-rescuing ClC-5 vector, we introduced the following mutations in the *CLCN5* gene: V523del (c.1566-1568del), E527D (c.1581A>T) and I524K (c.1571T>A). In addition, an HA-tag was also added in the C-terminus of each cDNA. Subsequently, all inserts were sub-cloned to an expression vector containing hygromycin resistance (pLenti CMV Hygro DEST 117-1, Addgene) using the Gateway recombination system. All constructs generated were stably transduced into previously *CLCN5* silenced cells using lentiviral particles produced in HEK-293 cells and were subsequently selected with 400 μg/ml hygromycin (Invivogen, #ant-hg-5). Site-directed mutagenesis was performed with the QuikChange^®^ II XL Site-Directed Mutagenesis Kit (Agilent Technologies) and primers were designed using the QuikChange Primer Design tool (Agilent Technologies).

### RNA extraction and qPCR

Total RNA was isolated from cells using TRIzol^®^ Reagent (#15596-026, Life Technologies) following the manufacturer’s protocol. cDNA was reverse-transcribed using the High-Capacity cDNA Reverse Transcription Kit (Applied Biosystems, #4387406). Endogenous and exogenous levels of *CLCN5* mRNA were measured using SYBR green probes (Applied Biosystems) and normalized against TBP using the following primers: Endogenous *CLCN5:* 5’-GGGATAGGCACCGAGAGAT-3’ and 5’-GGTTAAACCAGAATCCCCCTGT-3’; Exogenous *CLCN5:* 5’-GGTTACACACAACGGGCGAT-3’, and 5’-CGTAATCTGGAACATCGTA-3’; and TBP: 5’-CGGCTGTTTAACTTCGCTTC-3’ and 5’-CAGACGCCAAGAAACAGTGA-3’. In order to validate the microarray analysis was used the following TaqMan probes (Applied Biosystems): STEAP1 (Hs00185180_m1), ZPLD1 (Hs00604192_m1), PTPRD (Hs00369913_m1), CDH1 (Hs01023895_m1), EMX2 (Hs00244574_m1), NR1H4 (Hs01026590_m1), EHF (Hs00171917_m1), TBP (Hs00427620_m1). Analysis was performed using the 7900HT Sequence Detection System (Applied Biosystems). Relative expression fold change was determined by the comparative 2^(-ΔΔCT)^ method after normalizing to TBP. For the analysis of XBP-1 splicing, we used the following primers: 5’-AAACAGAGTAGCAGCGCAGACTGC-3’ and 5’-TCCTTCTGGGTAGACCTCTGGGAG-3’.

### Western blot

Cells were lysed in SET buffer (10 mM Tris-HCl pH 7.4, 150 mM NaCl, 1 mM EDTA and 1% SDS) and the protein concentration was quantified by the BCA assay (Thermo Fisher Scientific, #23225). Equal amount of whole cell extracts were resolved by SDS-PAGE and transferred to PVDF membranes (Millipore, #ISEQ00010). Membranes were blocked with 5% non-fat dry milk diluted in PBS-T (PBS 1x, Tween-20 0.1%) for 1 hour and incubated overnight at 4 °C with the appropriated antibodies: HA (dilution 1:1000, Roche, #867423001), β-tubulin (dilution 1:5000, Sigma, #T4026), E-Cadherin (dilution 1:1000, BD Transduction Labs, #610181), PERK (dilution 1:1000, Cell Signaling, #5683T), Cytokeratin-7 (Ventana Medical Systems, #790-4462) and Cytokeratin-18 (dilution 1:1000, Santa Cruz, #51582). Membranes were then incubated with the corresponding secondary antibodies (rabbit anti-mouse IgG/HRP, Dako, #P0260 and goat anti-rat IgG/HRP, Sigma, #A9037) at a 1:5000 dilution. Membranes were visualized using chemiluminescence reagent (Millipore, #WBLUF0500) and exposed on Odyssey Fc Imaging System (Li-Cor).

### Immunocytochemistry (ICC)

RPTEC/TERT1 cells were cultured on glass coverslips (Marlenfeld GmbH & Co. KG) for 10 days. Cells were then washed in cold PBS and fixed in −20□°C methanol for 5□min at room temperature. Aldehyde groups were quenched in 50 mM NH_4_Cl/PBS for 30 min and non-specific binding sites were blocked with 5% BSA in PBS for 60 min. Coverslips were incubated overnight at 4 °C with a 1:100 dilution with one of the following primary antibodies: HA (Roche, #867423001), KDEL (Abcam, #Ab10C3), Rab 5 (Cell Signaling, #3547S) and N-Cadherin (BD Transduction Labs, #610920), followed by incubation with the corresponding fluorescent-conjugated secondary antibodies (1:500 dilution, AlexaFluor Thermo Fisher Scientific #A11004, #A28175, #A11011, #A27034, #A21247, #A27012) for 1h at room temperature. Finally, cells were incubated with Hoechst 33342 (1:2000 dilution) (Invitrogen, #H1399) for 5 min to stain cell nuclei. Coverslips were then mounted on slides with Prolong diamond mounting medium (Thermo Fisher Scientific, #P36961) and fluorescence labeling was visualized in a confocal spectral Zeiss LSM 980 microscope. Acquired images were processed using ImageJ software and the co-localization analysis was performed using the ImageJ JACoP plugin. Co-localization was measured using Pearson’s and Manders co-localization coefficients.

### Glycosylation assay

RPTEC/TERT1 cells were lysed using RIPA buffer supplemented with protease inhibitor cocktail (Sigma-Aldrich, #P8340) and equal amounts of whole cell extracts were digested for 18 hours with endoglycosidase H (Endo H, New England, #P0702S) or peptide N-glycosidase F (PNGase F, New England #P0704S) enzymes following the manufacturer’s instructions.

### Albumin uptake

Albumin uptake was measured to investigate receptor-mediated endocytosis. RPTEC/TERT1 cells were seeded on glass coverslips (Marlenfeld GmbH & Co. KG) and grown for 10 days. To measure albumin uptake, cells were exposed to 50 μg/mL Alexa Fluor 488-conjugated Albumin (Thermo Fisher Scientific, #A13100) for 60 min. At the end of the incubation period, cells were washed 6 times with ice-cold PBS and fixed in −20□°C methanol for 5□min. To delimitate the cellular perimeter, slides were incubated overnight at 4 °C with a 1:100 dilution of N-cadherin antibody (BD Transduction Labs, #610920) followed by incubation with secondary fluorescence antibody Alexa Fluor Thermo Fisher Scientific, #A21247) for 1h at room temperature. Cell nuclei were stained with Hoechst 33342 (1:2000 dilution) (Invitrogen, #H1399) for 5 min at room temperature and slides were mounted with Prolong diamond mounting medium (Thermo Fisher Scientific, #P36961). Images were acquired with a confocal laser scanning microscope (Zeiss LSM 980) and processed using ImageJ software. Co-localization analysis was performed using the ImageJ JACoP plugin.

### DNA Microarray

Total RNA for DNA microarray was isolated as indicated before. RNA quality was checked using Bioanalyzer nano assay (Agilent Technologies). RNA samples representing five separate experiments from each of the conditions were used. Ten independent microarrays were performed using the Clariom D arrays (Affymetrix-Genechip array, #902922) according the manufacturer’s protocol.

### Over representation analysis

GO terms over-representation analysis was performed using the webserver g:Profiler (https://biit.cs.ut.ee/gprofiler/gost) as described in [46].

### Cell proliferation

Cell proliferation was performed as previously described [47]. Briefly, cells were incubated with 5 μM carboxyfluorescein succinimidyl ester (CFSE, Sigma-Aldrich, #21888) for 10 min at 37 °C. The unbound CFSE was quenched by washing cells twice in complete medium. An aliquot of cells was used to measure cell fluorescence at the onset of the experiment. The rest of labeled cells were seeded on tissue plates and incubated at 37 °C for 3 days. At the end of this period, fluorescence of daughter cells was measured. Cell fluorescence was measured on a FACS calibur flow cytometer (Becton Dickinson) and proliferation indices were determined using the Cell Quest software (Becton Dickenson).

### Cell adhesion

RPTEC/TERT1 cells cultured for 10 days were trypsinized, washed twice with culture medium to eliminate trypsin and counted. Fifty thousand cells/well were then seeded onto two duplicated 96-well plates for 60 minutes at 37°C. After this period, unattached cells from one of the plates were removed by washing cells twice with PBS, followed by an additional incubation period of 60 minutes in medium to facilitate cell recovery. The amount of attached cells (from the washed plate) and the total cells (from the unwashed plate) was determined using XTT assay (Sigma, # 11465015001) following the manufacturer’s instructions.

### Wound migration assay

For wound migration assay, 2.25 × 10^4^ cells were seeded on each of the two compartments of silicone culture inserts (Ibidi, #81176) and grown for 10 days. At the onset of the experiment, the culture insert was removed and cells were washed twice with medium to remove cell debris. Digital images were obtained every 30 minutes with a Thunder microscope (Leica) and area measurements were performed using ImageJ software.

### Statistical analysis

All statistical analyses were performed using GraphPad Prism 6 (RRID:SCR_002798) or SigmaPlot 10 (RRID: SCR_003210) software. Values are expressed as mean ± SEM. Statistical significance was determined by Student’s t test or one-way analysis of variance (ANOVA) followed by Turkey’s post hoc test. Criteria for a significant statistically significant difference were: *, p < 0.05; **, p < 0.01. Each specific test is indicated in figure legends.

---

**Figure S1.**
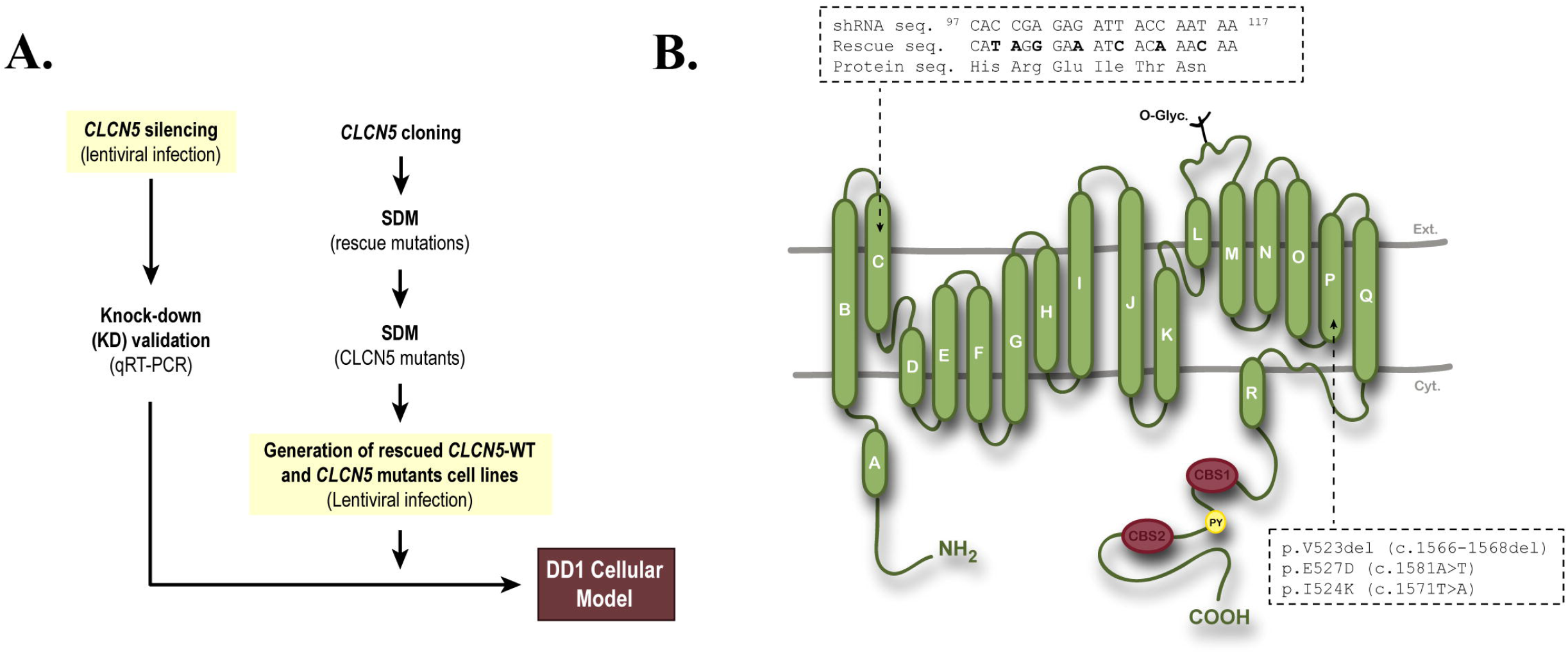
Generation of Dent disease 1 cell models. To explore the molecular mechanisms underlying PTCs dysfunction in DD1, we have generated stable RPTEC/TERT1 cell lines silenced for *CLCN5* gene or carrying the pathogenic ClC-5 mutations V523del, E527D or I524K. (A) CLCN5 was initially silenced in RPTEC(TERT1 cells using lentiviral shRNA vectors, and cells carrying *CLCN5* silencing were selected with the antibiotic puromycin. To re-introduce wild-type (WT) or mutant ClC-5, we introduced silent mutations in the shRNA target sequence to prevent RISC-mediated degradation. Subsequently, we mutated ClC-5 residues V523, E527 and I524K and we transduced the previously ClC-5 silenced cells. Cells carrying both CLCN5 shRNA and re-introduced ClC-5 forms were isolated using dual antibiotic selection (puromycin and hygromycin). (B) Scheme depicting ClC-5 the canonical 746-amino acid ClC-5 protein with its 18 membrane spanning α-helices, and the localization of shRNA target sequences and mutations V523, E527D and I524K within the helix P of ClC-5.

**Figure S2.**
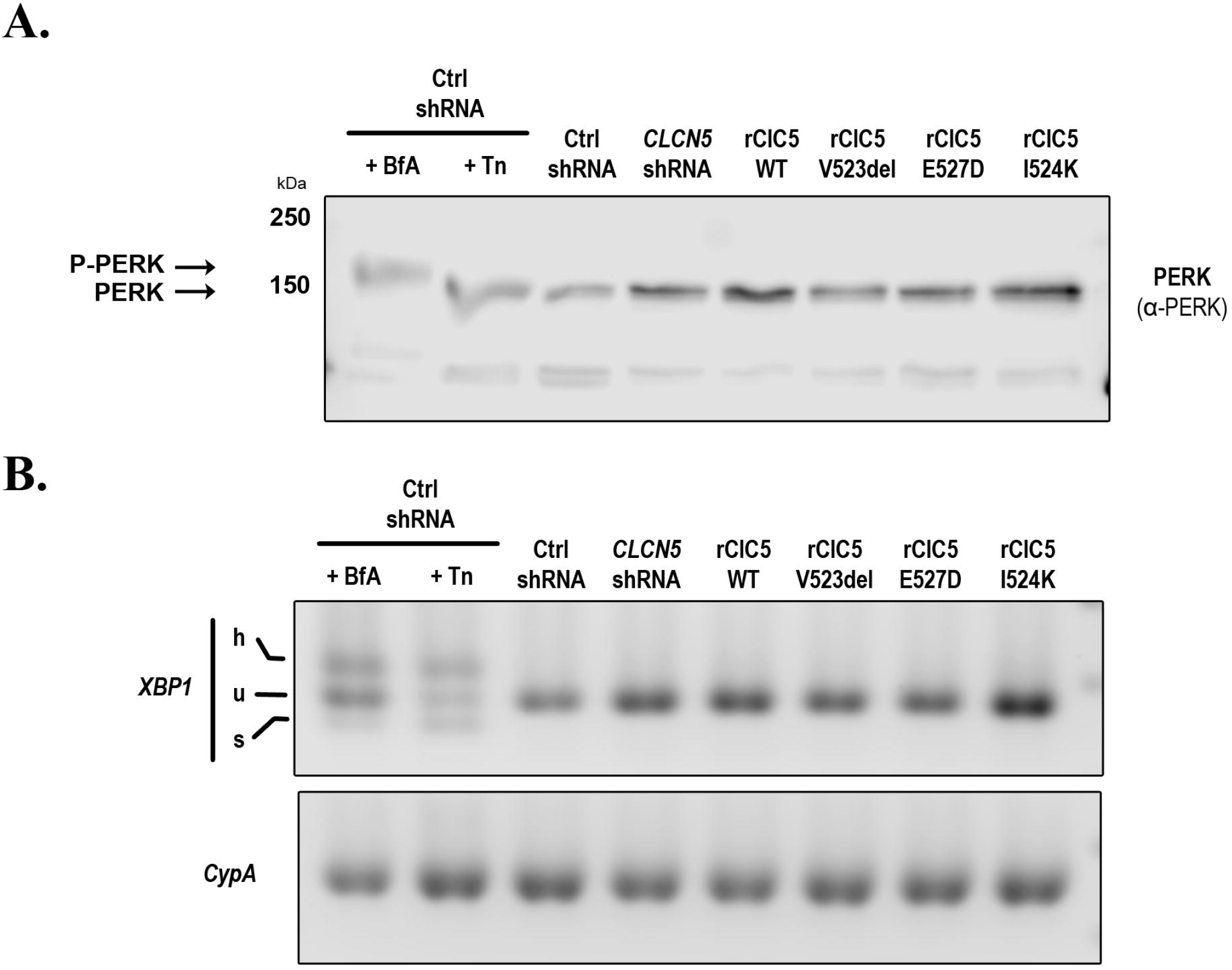
Expression of mutant ClC-5 proteins does not induce ER stress in RPTEC/TERT1 cells. To investigate whether the expression of ClC-5 mutants could be inducing the Unfolded Protein Response (UPR) and ER stress as a result of their accumulation in the ER, we checked the activation status of markers of the main branches of ER stress, i.e, PERK and XBP-1. As positive controls, cells were treated with the well-established ER stress inducers brefeldin-A (BfA) or Tunicamycin (Tn). (A) Western blots showing that only BfA, but not Tn, *CLCN5* silencing or expression of ClC-5 mutants induced a shift in the molecular weight of PERK, which has been associated with increased phosphorylation and activation of this protein kinase. (B) XBP-1 specific primers were used to analyze XBP-1 mRNA cleavage (u refers to unspliced and s to spliced mRNA) in cells expressing WT or mutant ClC-5. In this case, both BfA and TN induced XBP-1 cleavage. On the other hand, neither *CLCN5* silencing nor any of the ClC-5 mutants induced detectable cleavage of XBP-1.

**Figure S3.**
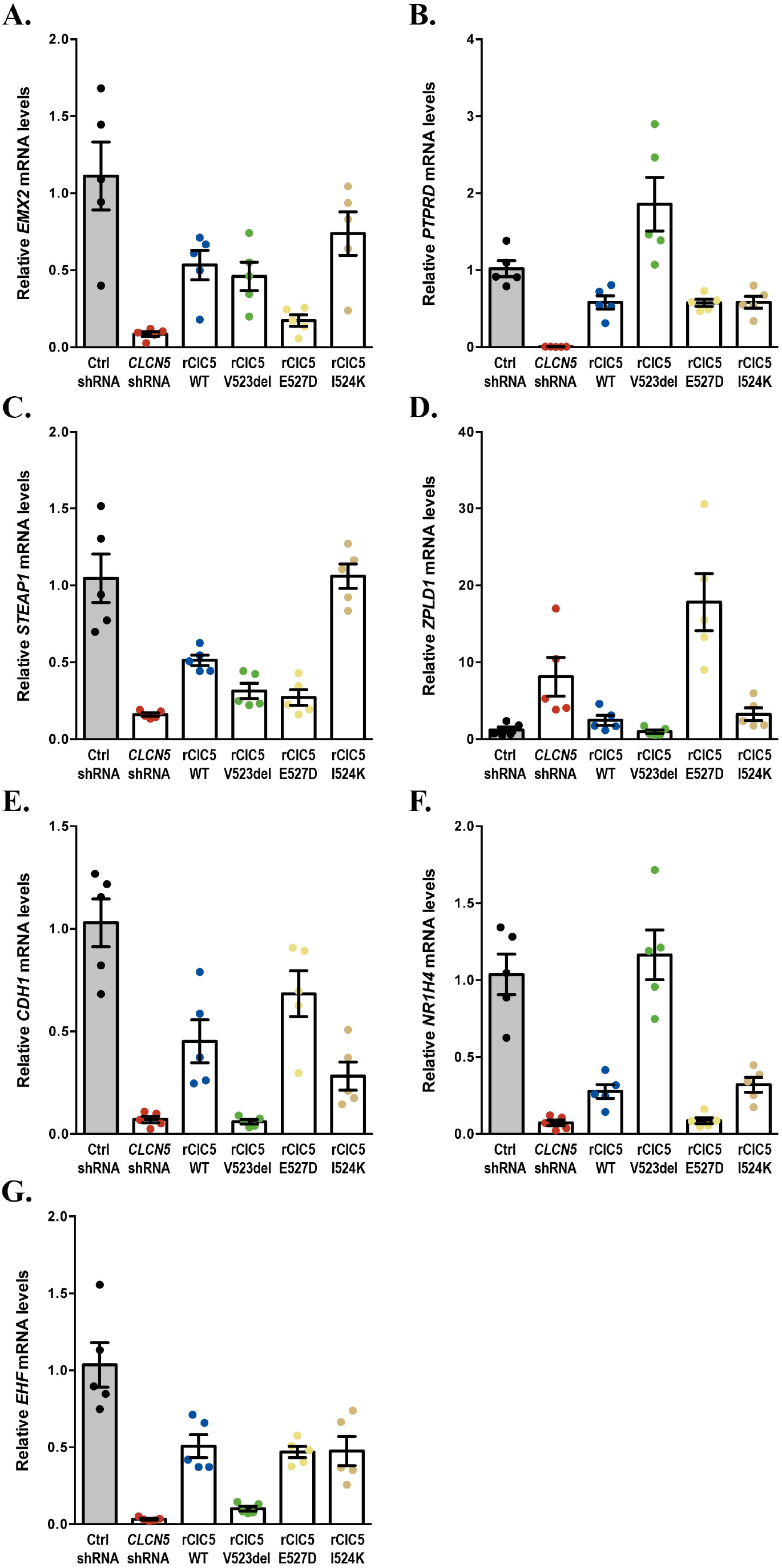
mRNA expression levels of DNA microarray validation genes. To validate the reliability of the results obtained from the DNA microarray, the expression levels of EMX2, PTPRD, STEAP1, ZPLD1, CDH1, NR1H4 and EHF genes were analyzed by qRT-PCR. Validation genes were selected among those that meet the following requirements: i) their expression was altered by some of the mutations compared to the WT condition and, ii) its expression was also modified by the silencing of *CLCN5* and totally or partially restored by ClC-5 WT re-introduction. All genes showed an expression pattern that correlated with that observed in the DNA microarray.

**Figure S4.**
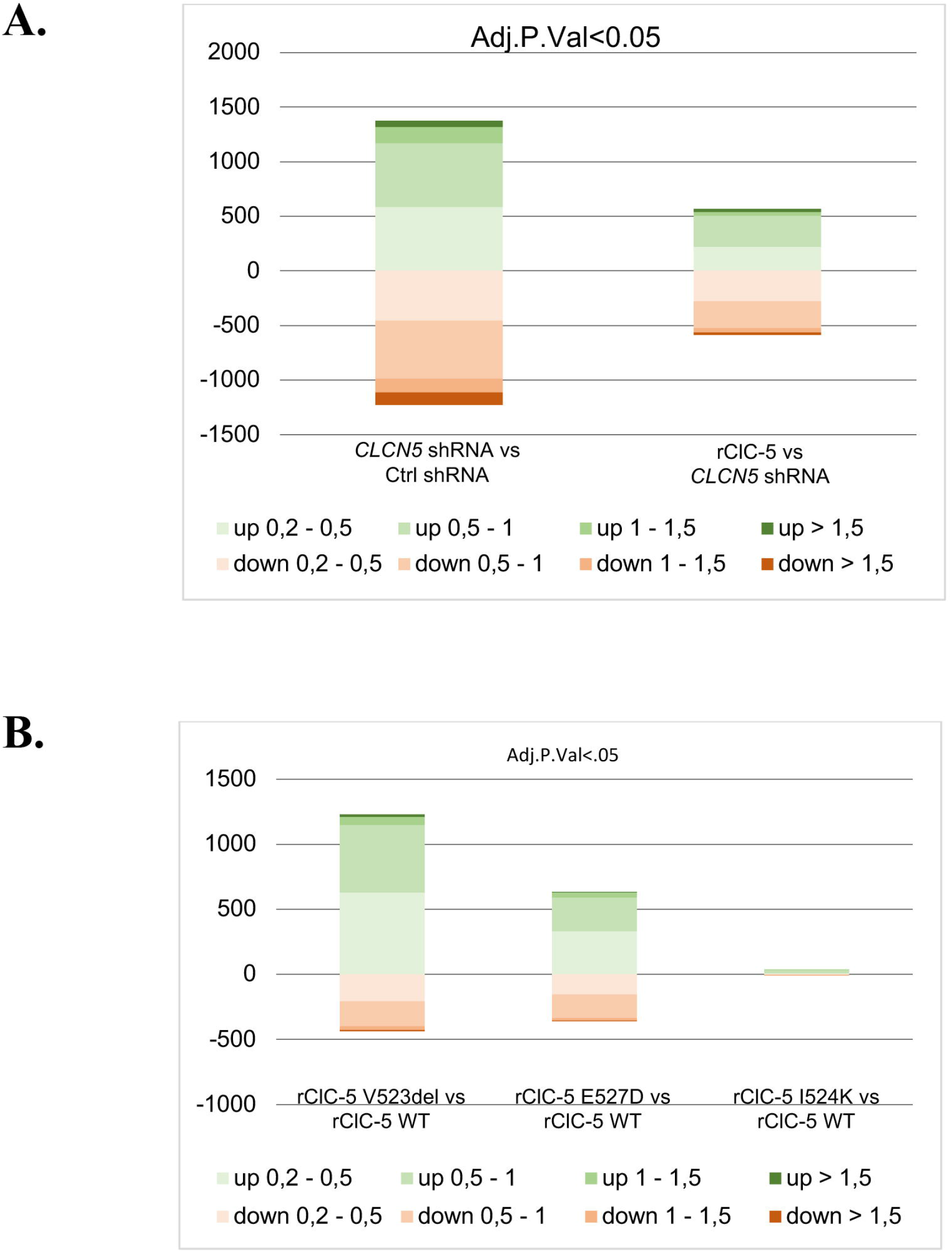
Number of genes in each range of differential expression. The number of genes in the DNA microarray whose expression was altered within a range of logFC for an adjusted p value lower than 0.05 in each of the comparisons that we have performed in this work.

