## Supplemental Tables for "Global gene-expression analysis reveals the molecular processes underlying ClC-5 loss-of-function in novel Dent Disease 1 cellular models"

Supplementary Tables

Table S1. Top 20 down-regulated genes by *CLCN5* silencing

| Gene Symbol | logFC.SHCLCN5vsSH0 | adjPval.SHCLCN5vsSH0 | logFC.WTvsSHCLCN5 | adjPval.WTvsSHCLCN5 |
| --- | --- | --- | --- | --- |
| SLPI | -5,012258677 | 7,25872E-18 | 2,490688757 | 1,48614E-10 |
| PTPRD | -4,774660354 | 5,73261E-18 | 4,076780087 | 4,04796E-16 |
| MUC1 | -4,725636323 | 1,63254E-20 | 2,641888478 | 1,76224E-14 |
| SOSTDC1 | -4,167052862 | 4,70927E-14 | 1,301344707 | 0,000249537 |
| EHF | -4,10574689 | 4,93531E-19 | 3,279045767 | 2,02258E-16 |
| SLC34A2 | -4,012080704 | 3,34897E-16 | 3,063994515 | 2,63198E-13 |
| ARHGEF38 | -3,797136664 | 7,14438E-15 | 0,983246057 | 0,000762179 |
| LIX1 | -3,7193965 | 9,11766E-14 | 0,883886992 | 0,005773676 |
| CDH1 | -3,697591042 | 7,14438E-15 | 2,466205116 | 1,48614E-10 |
| CXCL6 | -3,547100112 | 5,63081E-12 | 2,069829383 | 7,08157E-07 |
| ANPEP | -3,497712417 | 2,57939E-14 | 3,384013567 | 1,46952E-13 |
| C3 | -3,431960618 | 7,14438E-15 | 1,977200085 | 3,2444E-09 |
| PTGFR | -3,339062198 | 4,36867E-13 | 0,641977911 | 0,040806607 |
| CDH3 | -3,31371677 | 5,73261E-18 | 0,515618763 | 0,004751952 |
| LOC101927630 | -3,312773819 | 3,67966E-14 | 1,846828299 | 2,36354E-08 |
| DPP4 | -3,30097477 | 1,0374E-17 | 2,391987241 | 4,87344E-14 |
| MAOA | -3,287033568 | 1,28465E-14 | 1,55734935 | 1,80241E-07 |
| CSGALNACT1 | -3,242946306 | 1,80999E-13 | 0,723010169 | 0,012523965 |
| KRT7 | -3,191449694 | 1,43739E-14 | 1,843985356 | 5,77037E-09 |
| EMP1 | -3,03942192 | 4,08616E-14 | 1,579039068 | 9,30668E-08 |

Table S2. Top 20 up-regulated genes by *CLCN5* silencing

| Gene Symbol | logFC.SHCLCN5vsSH0 | adjPval.SHCLCN5vsSH0 | logFC.WTvsSHCLCN5 | adjPval.WTvsSHCLCN5 |
| --- | --- | --- | --- | --- |
| SLC14A1 | 5,08104134 | 4,79114E-14 | -3,977038387 | 4,18868E-11 |
| TTC29 | 3,679206623 | 1,14704E-12 | -1,067780535 | 0,002680413 |
| TMEM178A | 3,350740312 | 3,79059E-16 | -2,614100467 | 2,12546E-13 |
| ACSM3 | 3,04935039 | 3,89094E-14 | -3,482219987 | 7,56065E-15 |
| DIRAS2 | 3,014635521 | 3,92929E-15 | -1,807712049 | 6,97955E-10 |
| RAB3B | 2,969127618 | 7,25872E-18 | -1,363054288 | 6,97955E-10 |
| LINC01508 | 2,871168794 | 5,11238E-11 | -1,560351427 | 1,15104E-05 |
| ADAMTS16 | 2,856340255 | 2,26768E-10 | -2,292388697 | 6,32122E-08 |
| ITGA11 | 2,820938391 | 1,54507E-14 | -2,796221655 | 5,08676E-14 |
| KIT | 2,784145733 | 5,37046E-11 | -1,970813304 | 1,67632E-07 |
| NTM | 2,766697018 | 4,97022E-13 | -1,45799613 | 5,89012E-07 |
| FAM189A1 | 2,617138339 | 2,24867E-15 | -1,498680451 | 1,08268E-09 |
| SLC16A10 | 2,541950339 | 3,98234E-11 | -1,243742469 | 4,16723E-05 |
| ST8SIA2 | 2,44775401 | 7,32502E-14 | -1,116925328 | 1,39984E-06 |
| PGM5 | 2,44469443 | 1,80999E-13 | -1,476028667 | 2,36354E-08 |
| ZPLD1 | 2,429899703 | 1,65216E-12 | -1,687245584 | 1,20979E-08 |
| MAP3K7CL | 2,357575992 | 2,55733E-11 | -1,568185814 | 2,48828E-07 |
| SFRP4 | 2,331536894 | 7,59206E-14 | -1,353862223 | 2,35953E-08 |
| HMCN1 | 2,26960979 | 1,77002E-10 | -2,34963613 | 4,42059E-10 |
| ZNF385D | 2,161982015 | 1,33712E-12 | -1,526276707 | 6,79872E-09 |

Table S3. Top 20 down-regulated genes by re-introduction of V523del ClC-5 mutant

| Gene Symbol | logFC.V523delvsWT | adjPval.V523delvsWT |
| --- | --- | --- |
| CDH1 | -2,2699238 | 2,1101E-09 |
| MFAP5 | -2,1515998 | 2,1095E-08 |
| C3 | -2,0811644 | 2,9962E-09 |
| LUM | -1,8057315 | 5,6068E-07 |
| KRT7 | -1,7459962 | 2,9746E-08 |
| OPRPN | -1,6387219 | 1,2899E-05 |
| EHF | -1,5793078 | 3,4844E-09 |
| WNT7A | -1,5521822 | 2,1101E-09 |
| SLCO2B1 | -1,4213937 | 4,7274E-06 |
| ADGRF1 | -1,4152315 | 1,2046E-09 |
| FABP3 | -1,3842962 | 4,9308E-06 |
| MAP1B | -1,3263065 | 8,4725E-07 |
| MUC1 | -1,3139873 | 7,0926E-08 |
| ENTPD1 | -1,3088421 | 8,5295E-06 |
| MACC1 | -1,296171 | 7,5965E-09 |
| LCN2 | -1,2896225 | 0,00088447 |
| TAGLN | -1,2727844 | 1,6752E-05 |
| UCA1 | -1,2380877 | 5,3125E-08 |
| PDE1C | -1,2030441 | 7,0256E-06 |
| ZDHHC15 | -1,1699512 | 3,5396E-05 |

Table S4. Top 20 up-regulated genes by re-introduction of V523del ClC-5 mutant

| Gene Symbol | logFC.V523delvsWT | adjPval.V523delvsWT |
| --- | --- | --- |
| SLC17A1 | 3,04935497 | 7,574E-09 |
| SLC17A3 | 2,81283694 | 1,4946E-08 |
| NR1H4 | 2,59213551 | 7,3806E-11 |
| CLDN2 | 2,44095847 | 3,5125E-07 |
| NR1D1 | 2,35073551 | 2,1101E-09 |
| HES1 | 2,31656296 | 5,3125E-08 |
| CXCL6 | 2,11838805 | 6,8979E-07 |
| APCDD1L-AS1 | 1,9331687 | 2,7107E-08 |
| RPL22L1 | 1,88554225 | 4,4577E-13 |
| SLC27A2 | 1,81289443 | 2,919E-07 |
| KCNJ15 | 1,77286807 | 0,00010917 |
| EFHB | 1,71213441 | 3,5743E-07 |
| LINC01291 | 1,70389896 | 1,4025E-07 |
| TCF4 | 1,67893034 | 2,8791E-08 |
| KLF10 | 1,6788553 | 1,941E-09 |
| MATN2 | 1,59817147 | 1,2046E-09 |
| PTPRD | 1,54984402 | 4,1356E-07 |
| PRKAR2B | 1,54571457 | 5,1072E-07 |
| ANPEP | 1,51082471 | 2,1612E-06 |
| RELN | 1,49928918 | 2,328E-05 |

Table S5. Top 20 down-regulated genes by re-introduction of E527D ClC-5 mutant

| Gene Symbol | logFC. E527DvsWT | adjPval. E527DvsWT |
| --- | --- | --- |
| MFAP5 | -2,3750789 | 8,8178E-09 |
| FLG | -2,0583374 | 5,6017E-09 |
| NETO1 | -1,8661723 | 4,6023E-06 |
| EMX2 | -1,5523818 | 7,3078E-08 |
| CPT1A | -1,5493772 | 2,704E-08 |
| LCP1 | -1,4606835 | 3,044E-05 |
| THY1 | -1,4440417 | 1,1259E-06 |
| C3 | -1,3513745 | 5,6331E-06 |
| ABCB5 | -1,3061963 | 0,00024186 |
| NR1H4 | -1,2872162 | 1,2428E-05 |
| NPAS2 | -1,2807504 | 1,4478E-09 |
| KHDRBS3 | -1,2806112 | 3,1426E-08 |
| SGIP1 | -1,1928716 | 1,8136E-07 |
| ADAMTS9 | -1,1748411 | 1,4828E-06 |
| CACNA2D3 | -1,1701907 | 1,0215E-07 |
| HS3ST2 | -1,1226335 | 0,00024107 |
| CLDN10 | -1,0989468 | 0,00021468 |
| GREM1 | -1,0892441 | 0,00012516 |
| MLPH | -1,0785273 | 5,6162E-08 |
| CDH13 | -1,0721972 | 0,00017979 |

Table S6. Top 20 up-regulated genes by re-introduction of E527D ClC-5 mutant

| Gene Symbol | logFC. E527DvsWT | adjPval. E527DvsWT |
| --- | --- | --- |
| ZPLD1 | 2,24788773 | 5,697E-10 |
| TNNT1 | 1,90773131 | 1,1518E-08 |
| HLA-DRB1 | 1,79629489 | 8,8178E-09 |
| SELENOP | 1,7741324 | 8,6929E-07 |
| CLDN2 | 1,69304491 | 0,00011843 |
| HLA-DMA | 1,55848284 | 7,3078E-08 |
| GPX3 | 1,52380486 | 3,3773E-06 |
| MAPRE3 | 1,51464658 | 2,8941E-06 |
| NDST3 | 1,49303626 | 5,3534E-07 |
| LINC01508 | 1,45530881 | 6,5544E-05 |
| HLA-DPA1 | 1,44850854 | 1,029E-06 |
| SOSTDC1 | 1,42354969 | 0,00014937 |
| CORO2A | 1,38018793 | 5,8778E-06 |
| PAQR5 | 1,3748296 | 0,00038225 |
| LUCAT1 | 1,31447518 | 3,0936E-06 |
| SLC17A1 | 1,3092692 | 0,00193727 |
| SNAI1 | 1,29662188 | 0,00063583 |
| COBL | 1,29648308 | 3,9939E-06 |
| EFCAB13 | 1,2842711 | 0,00010262 |
| SPNS2 | 1,25758919 | 5,6162E-08 |

Table S7. Top 20 down-regulated genes by re-introduction of I524K ClC-5 mutant

| Gene Symbol | logFC.V523delvsWT | adjPval.V523delvsWT |
| --- | --- | --- |
| CDH1 | -1,10357 | 0,022233 |
| KRT7 | -0,74643 | 0,044569 |
| CATSPER1 | -0,66144 | 0,022233 |
| CLYBL-AS2 | -0,60521 | 0,044569 |
| SLC38A8 | -0,58758 | 0,044569 |
| CNGA2 | -0,51673 | 0,022233 |

Table S8. Top 20 up-regulated genes by re-introduction of I524K ClC-5 mutant

| Gene Symbol | logFC.I524KvsWT | adjPval.I524KvsWT |
| --- | --- | --- |
| PREX1 | 1,05644 | 0,022233 |
| CP | 1,031158 | 0,022233 |
| TMEM71 | 0,902159 | 0,022233 |
| STEAP1 | 0,752778 | 0,022233 |
| EXOSC8 | 0,70184 | 0,024853 |
| TCIM | 0,662846 | 0,035405 |
| IQCD | 0,660759 | 0,024853 |
| DYNC2H1 | 0,660272 | 0,02306 |
| LIN54 | 0,655919 | 0,023526 |
| LRTOMT | 0,629516 | 0,022233 |
| ZRANB3 | 0,628185 | 0,024853 |
| PLAG1 | 0,624192 | 0,022233 |
| TNIK | 0,614592 | 0,023526 |
| TEX2 | 0,58021 | 0,024853 |
| MIS12 | 0,572103 | 0,022233 |
| STEAP1B | 0,567719 | 0,044569 |
| NUFIP1 | 0,558783 | 0,022233 |
| PXYLP1 | 0,555539 | 0,044569 |
| CHCHD7 | 0,554923 | 0,022233 |
| RAB38 | 0,55239 | 0,044569 |
